# Polycomb Repressive Complex 2 is critical for mouse cortical glutamatergic neuron development

**DOI:** 10.1101/2023.11.05.565413

**Authors:** Laura Currey, Benjamin Mitchell, Majd Al-Kahlily, Sarah-Jayne McElnea, Danyon Harkins, Alexandra Pelenyi, Nyoman D. Kurniawan, Thomas H. Burne, Lachlan Harris, Stefan Thor, Michael Piper

## Abstract

The Polycomb Repressive Complex 2 (PRC2) regulates corticogenesis, yet the consequences of mutations to this epigenetic modifier in the mature brain are poorly defined. Importantly, PRC2 core genes are haploinsufficient and causative for several human neurodevelopmental disorders. To address the role of PRC2 in mature cortical structure and function, we conditionally mutated the PRC2 gene *Eed* in the developing mouse telencephalon. Adult homozygotes displayed smaller forebrain structures. Single-nucleus transcriptomics revealed glutamatergic neurons were particularly affected, exhibiting dysregulation of cortical layer-specific profiles, accompanied by aberrations in neuronal morphology and connectivity. Incredibly, homozygous mice performed well on challenging cognitive tasks. In contrast, heterozygous mice did not exhibit anatomical or behavioral phenotypes but displayed dysregulation of neuronal genes and altered neuronal morphology that was strikingly different from homozygous phenotypes. Collectively, we demonstrate how alterations to PRC2 function shape the mature brain and reveal a dose-specific role for PRC2 in determining glutamatergic neuron identity.

## Introduction

The mouse cerebral cortex exhibits tremendous cell diversity and elaborate networks of connectivity that are essential for higher-order behavioural and cognitive tasks. Cortical development is driven by the coordinated proliferation, then differentiation of neural progenitor cells (NPCs)^1^. This begins with the symmetric proliferative division of neuroepithelial cells within the cortical ventricular zone. As development proceeds, neuroepithelial cells give rise to radial glial cells (RGC) that divide asymmetrically, producing a RGC daughter cell, and either a neuron (direct neurogenesis; early in corticogenesis) or a basal progenitor cell (BP) (indirect neurogenesis; later in corticogenesis)^1^. During late gestation, a subset of RGCs switch to producing glial cells, such as astrocytes. Precise regulation of the number of neuroepithelial cell and RGC divisions, and of the extent of direct versus indirect neurogenesis, is critical for producing the correct number of neurons and glia in the mature cerebral cortex^1^.

Epigenetic modifiers play a key role in the developing brain, by regulating NPC proliferation and differentiation, as well as coordinating post-mitotic cellular identity. One epigenetic modifier that plays a critical role in regulating NPC behaviour is the Polycomb Repressive Complex 2 (PRC2). PRC2 is responsible for catalysing mono-, di-, and tri-methylation of lysine 27 (H3K27me1/2/3), primarily on histone H3.3 and to a lesser extent on H3.1 and H3.2^2^. H3K27me3 attracts Polycomb Repressive Complex 1 (PRC1), which catalyzes ubiquitylation of lysine 119 of Histone H2A (H2AK119ub), culminating in transcriptional silencing^3^. The core components of PRC2 include Enhancer of Zeste Homolog 1/2 (EZH1/2), Suppressor of Zeste 12 (SUZ12), Retinoblastoma binding protein 4/7 (RBBP4/7), and Embryonic ectoderm development (EED). Loss-of-function in any of these core components results in a lack of PRC2 function. Importantly, as PRC2 is the only known complex to catalyse H3K27me1/2/3, its function is absolutely essential to establish this repressive mark^4^, and so to mediate pivotal processes including ESC differentiation^5–7^, anteroposterior axis specification^8^, osteogenesis^9^ and neurogenesis^10^.

PRC2 function is critical in humans; the gnomAD and EXAC databases show *EED*, *EZH2* and *SUZ12* to be fully intolerant to a loss of gene dosage, and hence are haploinsufficient (Karczewski et al., 2020; Lek et al., 2016). Moreover, Weaver, Weaver-like, Cohen-Gibson syndromes, all of which are overgrowth disorders, are linked to heterozygous, likely neomorphic or hypermorphic gain-of-function mutations to *EED*, *EZH2* or *SUZ12*^11–13^. However, despite the clinical significance of these syndromes, our understanding of how PRC2 haploinsufficiency within the nervous system leads to these clinical outcomes is limited.

In mice, constitutive knockout of PRC2 core components causes gastrulation defects and embryonic lethality^14–18^. Conditional knockout of *Eed* in the early CNS using a *Sox1-Cre* driver [expressed in all NPCs from embryonic day (E) 8.5], culminates in embryonic microcephaly arising from premature NPC differentiation^19,20^. Likewise, conditional deletion of *Ezh2* from dorsal telencephalic NPCs at E9.5 causes early onset of neurogenesis at the expense of neuroepithelial cell self-renewal, resulting in NPC depletion and consequently fewer late-born neurons^21^, a phenotype mirrored by the deletion of *Eed* from dorsal telencephalic NPCs^22^. Collectively, these studies point to precocious NPC differentiation as a key phenotype arising from abnormal PRC2 function early in CNS development. Numerous other genetic perturbations also cause premature NPC differentiation^23–28^. However, the adult phenotypes in these mice, including those with abnormal PRC2 function, has received limited attention. PRC2 function also contributes to neuronal specification. In the mouse hypothalamus, loss of *Eed* has been reported to have subtle effects on cellular specification, including a reduction in dopamine-, Hypocretin-, and Tac2-Pax6-expressing neurons, and an increased number of neurons expressing both glutamatergic and GABAergic markers^29^. PRC2 is also involved in the survival and function of adult striatal neurons^30^, and in maintaining identity and function of dopaminergic and serotonergic neurons^31^. However, whether the loss of *Eed* impacts neuronal specification and identity within the cerebral cortex remains unclear.

Here, we sought to address how precocious NPC differentiation manifests within the adult brain, and whether PRC2 function regulates neuronal identity in the cortex, using mice in which *Eed* had been removed from all cells derived from the dorsal telencephalon (*Emx1-iCre*). Adult homozygous (*Eed-cKO)* but not heterozygous (*Eed-cHet*) knockout mice exhibited microcephaly. Notably, the absence of *Eed* resulted in loss of identity in glutamatergic neurons including downregulation of neuron-specific genes, abnormal expression of non-neuronal genes and aberrant laminar identity. In contrast, there were minimal disruptions to gene expression in other cells including astrocytes and oligodendrocytes. To our surprise, *Eed-cKO* mice displayed remarkably normal behaviour across a range of cortically relevant behavioural tests, some of which were cognitively demanding. Collectively, our findings elucidate the cellular, structural, and transcriptomic deficits arising from PRC2 loss-of-function and identifies heretofore unrealised plasticity with regards to how these mice complete complex behavioural tasks.

## Results

### Loss of *Eed* results in a smaller cortex in the adult

Conditional ablation of PRC2 genes from CNS progenitor cells has been shown to cause severe reduction of cortical size in the embryo^19,21,22^. How does this phenotype manifest in the adult brain? To investigate this question we employed a *Eed-Emx1-iCre* model, in which *Eed* is conditionally removed from all cells derived from the dorsal telencephalon NPCs, including glutamatergic neurons, astrocytes, and most cortical oligodendrocytes, but not cell types which arise from other regions during development, such as interneurons, microglia and a proportion of oligodendrocytes^32,33^. Consistent with previous embryonic data, we found that *Eed-cKO* brains had major reductions in tissue size throughout cortical and hippocampal regions (Figure 1, Supplementary 1). By contrast, there was no significant difference between control and *Eed-cHet* brains. The width of the *Eed-cKO* cortical plate was reduced in the cingulate, motor, sensory, auditory, and visual cortices (Figure 1G, H). The retrosplenial cortex was also reduced in more caudal sections (Supplementary 1Q).

**Figure 1:**
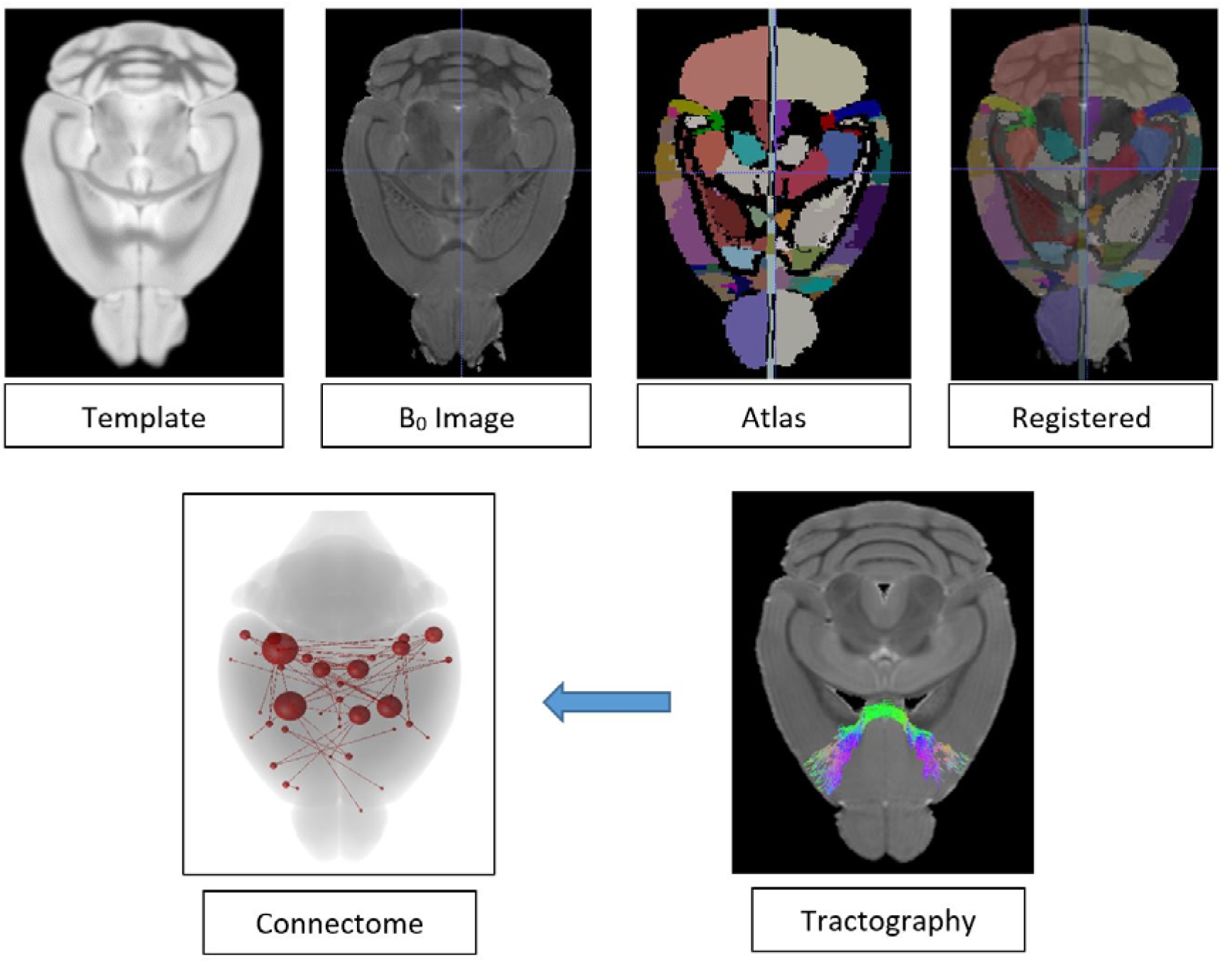
Generation of structural connectome.

The dramatic reduction in size of the dorsal telencephalon of cKO mice prompted us to examine if the three major forebrain commissures were intact, given they arise, in part, from tissue derived from Emx1-expressing progenitor cells. The corpus callosum of *Eed-cKO* mice was severely diminished, and callosal axons were rarely observed to cross the midline (Figure 1C). The only region we observed *Eed-cKO* callosal axons decussating was at approximately Bregma 0.85mm (Supplementary 1C), at which point the dorsoventral height of the corpus callosum at the midline measured just 7% of the average control. The hippocampal commissure was difficult to identify in *Eed-cKO* hematoxylin-stained sections (Figure 1C). This was likely due to the hippocampal phenotype, with the dentate gyrus and the CA1-3 regions either severely reduced or unidentifiable (Figure 1F). The penetrance was 100%, with all animals analysed showing cortical defects. By contrast, expressivity was variable, particularly in the hippocampus where phenotypes ranged from a complete absence of hippocampal structures to severely diminished, but present, CA or dentate gyrus regions (Supplementary 2E-T). There was no clear correlation between gender and severity of phenotype. In contrast to the corpus callosum and hippocampal commissure, the anterior commissure was not significantly altered in either heterozygous or homozygous animals.

We next investigated how the reduction in cortical size manifested at a more global level with high-resolution episcopic microscopy (HREM), a technique that enables 3D volume rendering to be performed upon embedded and sectioned tissues (Figure 1I-S). Consistent with our hematoxylin analyses, *Eed-cKO* brains were severely reduced in volume. On average, the whole brain volume (excluding the olfactory bulbs and cerebellum) was 43% smaller in *Eed-cKO* brains compared to controls. In line with the hematoxylin results, the volume of the neocortex, hippocampus, and hippocampal commissure (including fornix and fimbria) was reduced in *Eed-cKO* mice. Although *Emx1* is expressed during the development of the olfactory bulb and in a proportion of olfactory cells in the adult^34,35^, there was no significant reduction in the volume of the olfactory bulb, olfactory tubercle, or olfactory cortex of *Eed-cKO* mice. The thalamus was also smaller in *Eed-cKO* mice, likely an indirect effect since this region is not deleted for *Eed*. We did not detect any differences between control and *Eed-cHet* brains, neither with regards to whole brain volume, nor with discrete regions of the telencephalon or diencephalon. Volumetric magnetic resonance imaging further confirmed the reduction in a variety of cortical and subcortical structures of adult *Eed-cKO* mice (Supplementary 2A-D). Collectively these findings indicate that removal of *Eed* from neural progenitor cells of the developing dorsal telencephalon culminates in a dramatic reduction in adult cortical size.

### Loss of *Eed* causes laminar deficits in the cortical plate

Premature differentiation of neural progenitor cells is known to contribute to microcephaly, and to preferentially affect late born neurons within the cerebral cortex^36^. To investigate lamination in our model, we performed immunofluorescent staining of the neuronal markers Satb2, Ctip2, and Foxp2 (Figure 2A-I). Satb2 is expressed in cortical layers 2-6, Ctip2 is expressed in layers 5/6, and Foxp2 is expressed in layer 6, allowing different cortical layers to be distinguished^37,38^.

**Figure 2.**
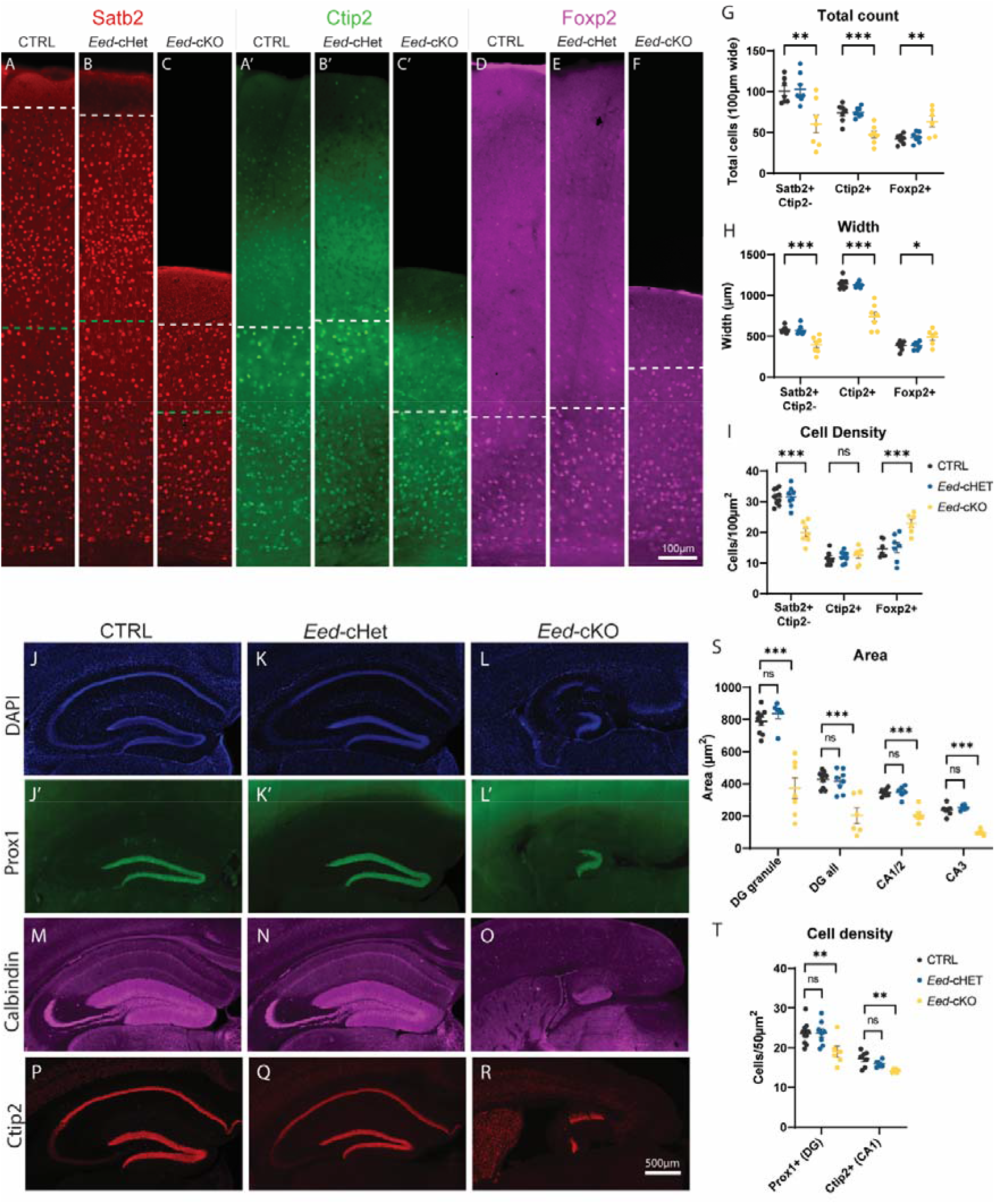
Loss of *Eed* causes deficits in cortical lamination and hippocampal morphology. (A-F) Immunofluorescence staining of Satb2 (A-C), Ctip2 (A’-C’), and Foxp2 (D-F) of the CTRL, *Eed-cHet*, and *Eed-cKO* somatosensory cortex. Satb2 and Ctip2 were co-stained, whereas Foxp2 was stained from separate sections. Dashed lines represent the border between positive and negative layers for each marker. (G) The total number of positive cells within a 100 µm wide region spanning all layers of the cortical plate was counted. (H) The width of Ctip2+, Ctip2-/Satb2+, and Satb2+ cortical layers was measured. (I) Cell density was obtained by counting number of cells within a 100 µm^2^ region within layer 6 (Foxp2), layer 4/5 (Ctip2), or layer 2/3 (Satb2). *** = p<0.001, ** = p<0.01, * = p<0.05, ns = not significant, one-way ANOVA. n = 7 CTRL, 7 *Eed-cHet*, 7 *Eed-cKO* (G-I). See also Supplementary 4 for additional images showing disrupted laminar organization. (J-R) Immunofluorescence staining of DAPI (J-L), Prox1 (J’-L’), Calbindin (M-O), and Ctip2 (P-R) in the hippocampus of CTRL, *Eed-cHet*, and *Eed-cKO* mice. DAPI and Prox1 were co-stained, whereas Calbindin and Ctip2 were stained from separate sections. (S) The area of the dentate gyrus granule layer (“DG granule”, indicated by Prox1+ staining), dentate gyrus region (“DG all”, granule, molecular, and polymorph layers combined, as indicated by Calbindin+ staining in the dentate gyrus region), CA1/2 pyramidal layer (Ctip2+) and CA3 pyramidal layer (Ctip-, DAPI+) were measured. (T) Cell density of Prox1+ cells in the dentate gyrus and Ctip2+ cells in the CA1 were measured. *** = p<0.001, ** = p<0.01, ns = not significant, one-way ANOVA. n = 10 CTRL, 8 *Eed-cHet*, 8 *Eed-cKO* (Prox1 area measurements); n = 10 CTRL, 8 *Eed-cHet*, 6 *Eed-cKO* (Prox1 cell count). n = 9 CTRL, 6 *Eed-cHet*, 6 *Eed-cKO* (Calbindin). n = 7 CTRL, 6 *Eed-cHet*, 6 *Eed-cKO* (Ctip2, DAPI). See also Supplementary 4 for width and length measurements of the dentate gyrus and CA1-3 regions.

Consistent with the hematoxylin stains, the *Eed-cKO* mice had a thinner cortical plate, with decreased width in both Satb2+/Ctip2- and Ctip2+ layers, denoting upper and mid-lower layers, respectively. In line with this, *Eed-cKO* mice also had fewer Satb2+/Ctip2- and Ctip2+ cells compared to controls. Remarkably however, the width of the Foxp2+ layer (i.e., layer 6) was thicker in *Eed-cKO* mice than controls, and had more Foxp2+ cells in total, despite the overall thinner cortical plate. Additionally, the cell density of Foxp2+ cells was also increased in *Eed-cKO* mice, whereas cell density of Satb2+/Ctip2-cells in layer 2-3 was decreased, and Ctip2+ cells in layer 4-5 was unchanged. There were no significant differences in *Eed-cHet* mice compared to controls. Interestingly, we also observed that the delineation between layers was abnormal in *Eed-cKO* mice. For instance, the boundary between Foxp2, Ctip2, of Satb2 positive and negative layers formed a distinct, straight line in *Eed-cHet* and control mice. However, in *Eed-cKO* mice, this boundary was erratic and often ambiguous (Supplementary 3A-I). Altogether, these findings indicate that *Eed-cKO* mice have substantial disorganisation of cortical lamination. The reductions in width and cell number in layers 2-5 is in line with microcephaly and precocious neurogenesis phenotypes. However, the increase in both width and cell number of layer 6 is unusual. One hypothesis arising from this is that neuronal specification is abnormal in the absence of H3K27me3.

Next, we investigated astrocytes and oligodendrocytes, via GFAP and Olig2 expression respectively (Supplementary 3U-AB). Interestingly, we observed an increase in GFAP+ expression in the motor/cingulate cortex of *Eed-cKO*, but not in other cortical regions. In contrast, density of Olig2+ cells was decreased in *Eed-cKO* mice, suggesting that the number of oligodendrocytes may be reduced. There was no difference in GFAP or Olig2 staining in *Eed-cHet* mice compared to control. Finally, we investigated interneurons with Calbindin immunofluorescent stains, which labels a subset of interneurons (Supplementary 3AC-AI). The density of Calbindin+ cells was increased in *Eed-cKO* mice. As interneurons are not affected by *Eed* deletion in this mouse model, this increased cell density is likely due to a normal number of interneurons migrating into an underdeveloped cortex.

### Loss of *Eed* causes altered hippocampal morphology

The dramatic hippocampal phenotype observed at a gross morphological level (Figure 1) led us to next investigate the hippocampus in more detail. To this end, three hippocampal markers were used - Prox1 to stain the granule layer of the dentate gyrus, Calbindin to stain the molecular, granule, and polymorphic layers of the dentate gyrus, and Ctip2 to stain the CA1/2 regions (Figure 2J-T). The CA3 region was inferred by DAPI+, Ctip2-staining. *Eed-cKO* sections were highly abnormal, often displaying a complete absence of Prox1+ cells in the presumptive dentate gyrus. Of the 10 *Eed-cKO* biological replicates examined, 3 samples did not exbibit any Prox1+ cells, 4 samples only had Prox1+ granule cells in one hemisphere, and 3 samples had a Prox1+ granule cells in both hemispheres. Similar results were also found for Calbindin and Ctip2 (Table 1). *Eed-cHet* samples did not exhibit observable differences compared to controls.

**Table 1:**
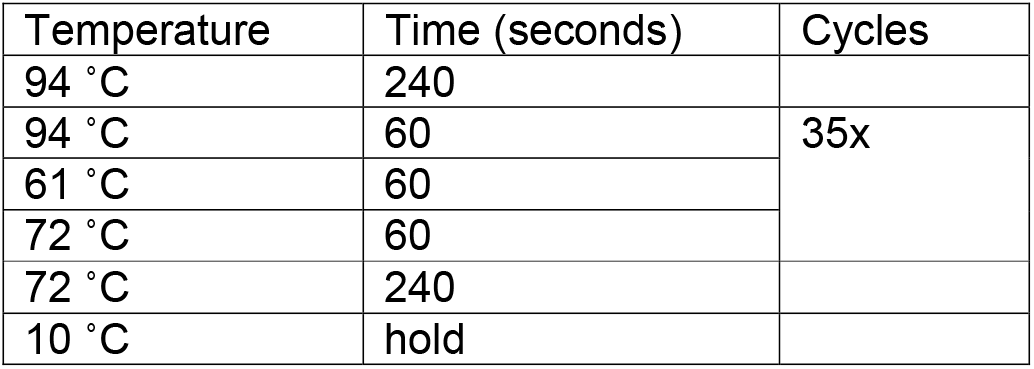
PCR cycle conditions.

For those sections that did show positive staining of Prox1, Calbindin, or Ctip2, the morphology of the dentate gyrus and CA regions was highly abnormal. *Eed-cKO* mice displayed substantial reduction in dentate gyrus area and in the density of Prox1+ granule cells. Furthermore, the length of the CA1/2 and CA3 regions were reduced in *Eed-cKO* mice. In contrast, the width of the CA1 region was increased in comparison to controls (Supplementary 3J-N). However, overall, the areas of both the CA1/2 and CA3 regions were smaller. Moreover, the density of Ctip2+ in CA1 region was reduced in *Eed-cKO* mice. Collectively, these results show that *Eed-cKO* mice have distinct hippocampal abnormalities and high phenotypic variability between samples and even between hemispheres.

### Glutamatergic neurons have aberrant laminar markers and ectopic gene expression in *Eed-cKO* mice

Our findings are consistent with previous studies linking PRC2 function to the neural stem cell proliferation and ultimately cortical size^19–22^. However, our lamination data hinted at previously unrecognised roles for this complex in mediating cortical glutamatergic cell identity. To interrogate this in more detail, we isolated nuclei from the neocortex of adult CTRL, *Eed-cHet* and *Eed-cKO* mice and performed single-nucleus RNA-sequencing (snRNA-seq). Following quality control, we sequenced and analysed 2,715, 4,660, and 3,755 high-quality sequenced nuclei from CTRL, *Eed-cHet*, and *Eed-cKO mice*, respectively (Figure 3). Uniform Manifold Approximation and Projection (UMAP) dimension reduction was used to visualize the data. Clustering of cells according to their broad characteristics revealed the expected cortical cell types [astrocytes, GABAergic neurons, glutamatergic neurons, microglia, oligodendrocytes, oligodendrocyte precursor cells (OPCs), and vascular cells] in all genotypes (Figure 3A). However, when we analyzed by genotype, an interesting pattern emerged. While CTRL, *Eed-cHet*, and *Eed-cKO* cells clustered together for most cell types (pre-oligodendrocytes, oligodendrocytes, astrocytes, microglia, and interneurons), we found that *Eed-cKO* glutamatergic neurons clustered separately from control and *Eed-cHet* cells (Figure 3B). This suggested that, while most cell types have high similarity in gene expression between genotypes, *Eed-cKO* glutamatergic neurons exhibit divergent gene expression.

**Figure 3.**
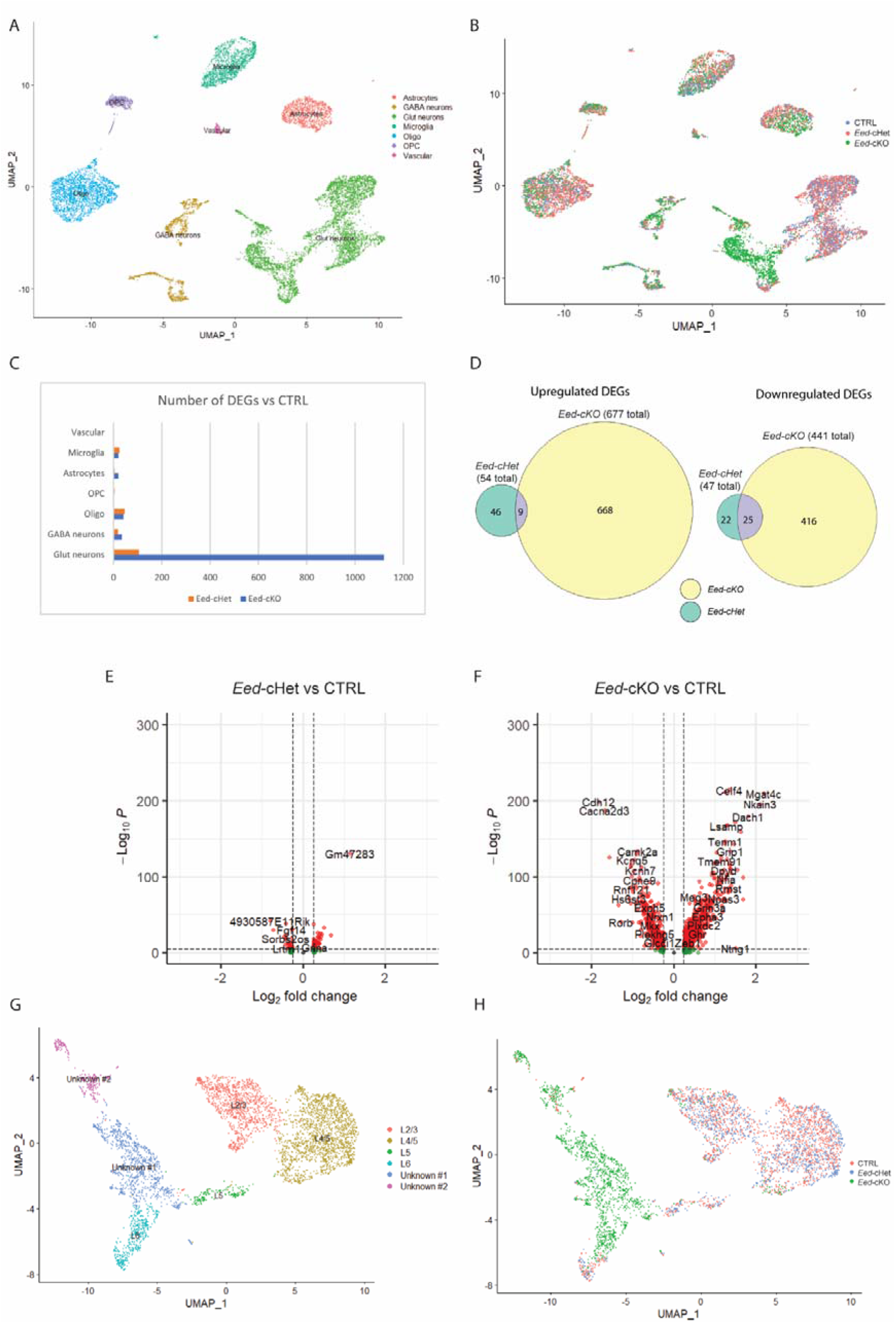
*Eed* regulates gene expression in glutamatergic neurons. (A) UMAP of combined cells from CTRL, *Eed-cHet*, and *Eed-cKO* samples, revealing that cells cluster by cell type. (B) UMAP of CTRL (red), *Eed-cHet* (blue), and *Eed-cKO* (green) cells, coloured by genotype to allow for intergenotype comparison. (C) Total number of differentially expressed genes (DEGs) for each major cell type in *Eed-cHet* and *Eed-cKO* mice compared to CTRL. (D) Overlap of DEGs common between both *Eed-cHet* and *Eed-cKO* for upregulated (left) and downregulated (right) genes. (E) Volcano plot showing DEGs in *Eed-cHet* and *Eed-cKO* glutamatergic neurons. (G,H) UMAP visualisation of glutamatergic neurons only. Cells are coloured according to cluster (G) or genotype (H), demonstrating that CTRL and *Eed-cHet* cells form clusters according to cortical layers, whereas *Eed-cKO* cells form separate clusters. Clusters which could not be categorised into cortical layers are labelled “Hom #1”, “Hom #2”, and “Hom #3”. Oligo = oligodendrocytes, OPC = oligodendrocyte progenitor cells, Glut = glutamatergic, micro = microglia.

This phenomenon was made clearer when we investigated the number of differentially expressed genes (DEGs) (Figure 3C). Interneurons and microglia are not deleted for *Eed* in this mouse model, and, as anticipated, we did not observe major alterations in gene expression in these cell types. However, astrocytes, a subset of oligodendrocytes, and a subset of pre-oligodendrocytes, which do arise from the dorsal telencephalon and are deleted for *Eed*, all displayed less than 50 DEGs each for both *Eed-cHet* and *Eed-cKO* samples, showing that gene expression was not substantially altered in these cell types. Collectively, these data suggests that although cortically derived glial cells lack *Eed* in this model, their gene expression profiles are relatively unaffected in cKO animals.

In stark contrast, glutamatergic neurons displayed substantially altered gene expression in *Eed-cKO* nuclei, with 1,119 DEGs (Figure 3C). Interestingly, despite the mild morphological and cellular effects observed above in the heterozygous animals (Figure 1, 2), *Eed-cHet* glutamatergic neurons displayed 105 DEGs (Figure 3C). When we compared the cKO versus the cHet DEGs, surprisingly, a relatively small percentage of DEGs were common between *Eed-cHet* and *Eed-cKO* glutamatergic neurons (Figure 3D). Approximately 16% of genes upregulated in *Eed-cHet* glut neurons were also upregulated in *Eed-cKO* glut neurons; and approximately 50% of genes downregulated in *Eed-cHet* glut neurons were also downregulated in *Eed-cKO* glut neurons. Next, glutamatergic neurons were segregated and re-clustered, allowing them to be investigated at greater resolution. By clustering these cells independently from other cell types, we anticipated that they would form further clusters, such as by cortical layer. Indeed, both control and *Eed-cHet* glutamatergic neurons formed clusters that could be classified into cortical layers (layers 2/3, 4/5, 5, and 6) based on canonical gene expression (Figure 3G-H). However, *Eed-cKO* samples did not form readily identifiable clusters based on laminar markers, as their expression of cortical markers was highly irregular. Instead, *Eed-cKO* glutamatergic neurons resolved into three abnormal clusters.

These data, in conjunction with our analysis of layer-specific markers (Figure 2), led us to posit that specification of glutamatergic neuronal identity may be aberrant in cKO animals, culminating in the abnormal expression of genes that could otherwise show distinct laminar specificity. Indeed, many of the top DEGs were genes that normally have cortical layer-specific expression. To investigate this further, we analysed specific candidate genes in our sequencing dataset. For example, *Foxp2* is usually expressed only in layer 6 projection neurons^39^. Control and *Eed-cHet* neurons had a single cluster strongly expressing *Foxp2*, indicating that these were layer 6 neurons. However, *Foxp2* was expressed throughout many *Eed-cKO* clusters. Similar results were found for other cortical layer markers, including *Lrrtm4* (L2/3), *Gpc6* (L2/3), and *Rorb* (L4/5) (Figure 4A-B). Consequently, *Eed-cKO* glutamatergic neurons could not be subcategorised. Not only were cortical layer markers spread across clusters, but they were also highly dysregulated in *Eed-cKO* neurons. For example, layer 2/3 genes *Lrrtm4* and *Rarb*, and layer 4 gene *Cdh12,* were upregulated. Many cortical layer genes were also downregulated, such as upper layer genes *Satb2*, *Cux1*, and *Pdzrn3*, and deep layer genes such as *Zmat4* and *Rorb* (Figure 4C). In addition to dysregulation of cortical layer markers, *Eed-cKO* neurons also showed ectopic expression of genes that are not normally expressed in glutamatergic neurons, including genes that are normally expressed in interneurons, glia, or in tissues outside of the CNS (Figure 4E-F). The combined effect of dysregulation of laminar-specific neuronal markers and ectopic gene expression suggests that loss of *Eed* compromises cell identity specifically in glutamatergic neurons.

**Figure 4.**
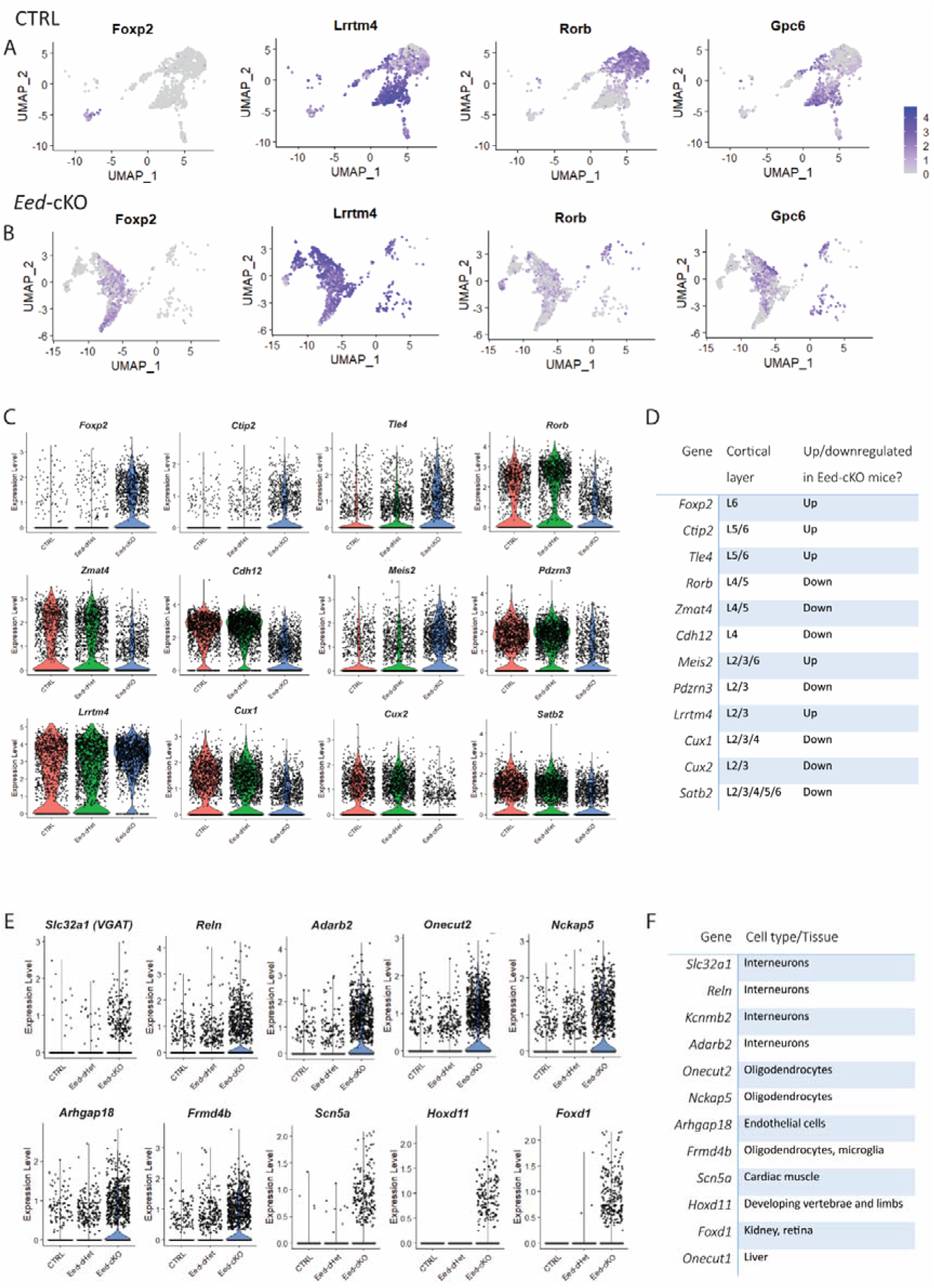
Glutamatergic neurons have aberrant laminar markers and ectopic gene expression in *Eed-cKO* mice. (A, B) UMAPs showing expression of cortical layer markers *Foxp2*, *Lrrtm4*, *Rorb*, and *Gpc6* (denoting layer 6, 4/5, 2/3, and 2/3, respectively) in CTRL (A) and *Eed-cKO* (B) glutamatergic neurons, allowing visualisation of how these markers are expressed across clusters. Dark blue represents cells with high expression of these genes. In CTRL cells, markers are largely expressed in a single cluster, whereas in *Eed-cKO* cells they are broadly expressed across clusters. (C) Violin plots showing examples of dysregulated gene expression of cortical layer markers in *Eed-cKO* neurons. (D) List of which cortical layers the genes in (C) are normally expressed in. Also listed is whether these genes were up or downregulated in *Eed-cKO* neurons. (E) Examples of genes that were ectopically expressed in *Eed-cKO* glutamatergic neurons. (F) A table showing examples of what cell types or tissues the genes in (E) are usually expressed in. See also Supplementary 4 for gene ontology analysis.

In line with this, gene ontology (GO) enrichment analysis identified that neuron-specific processes were highly implicated in *Eed*-cHet and *Eed*-cKO mice (Supplementary figure 7). In *Eed*-cKO neurons, top enriched biological process included terms such as synaptic organisation, cell junction assembly, neurotransmitter transition, membrane potential, transmembrane transport, axonogeneis and dendrite development. Top enriched molecular functions involved transmembrane transport, ion channels, and actin binding, and the synapse was the top implicated cell compartment. Eed-cHet neurons shared similar enriched GO terms. These data are consistent with the deficits in glutamatergic gene expression, further implicating *Eed* in the regulation of glutamatergic neuron identity.

### *Eed-cKO* pyramidal neurons have morphological deficits

Considering the enrichment of GO terms associated with axonogenesis and dendrite development, we sought to determine whether the morphology of glutamatergic neurons was abnormal in *Eed-cHet* and *Eed-cKO* mice. To investigate this, we used Golgi-Cox staining of pyramidal neurons in layers 2/3 and 5 of the motor cortex (Figure 5A). Interestingly, *Eed-cKO* neurons in layer 5 had reduced dendrite length (Figure 5B-C) and arborization (Figure 5B-C). By contrast, dendritic length and arborization was not altered in layer 2/3 *Eed-cKO* neurons. These changes may result from cell-intrinsic dendritic development defects or from the smaller cortex of *Eed-cKO* mice, as dendrites are unable to project their normal distances in the reduced tissue size. Remarkably, *Eed-cHet* neurons had a distinct phenotype from *Eed-cKO* neurons. Although layer 5 basal dendrites showed decreased arborization, akin to *Eed-cKO* neurons (Figure 5H), *Eed-cHet* neurons in layer 2/3 surprisingly had an increased number of branches in basal dendrites (Figure 5F). Next, we analyzed dendritic spines in layer 2/3 pyramidal neurons (Figure 5I-M). Spine density was not significantly altered in *Eed-cHet* or *Eed-cKO* mice compared to controls (Figure 5J). However, on average *Eed-cKO* dendritic spines had reduced length and *Eed-cHet* dendritic spines had increased head diameter (Figure 5K-L). Spines were further classified as thin, stubby, or mushroom spines depending upon their size and shape. In general, mushroom spines are mature with strong synaptic connections, thin spines are less mature but more dynamic, and stubby spines are thought to be immature, transitory structures^40^. We found that *Eed-cKO* mice had a lower proportion of mushroom spines (Figure 5S), whereas *Eed-cHet* mice had a lower proportion of thin spines, and a higher proportion of stubby spines (Figure 5M). Overall, these results suggests that *Eed-cHet* and *Eed-cKO* neurons have different deficits in spine maturation.

**Figure 5.**
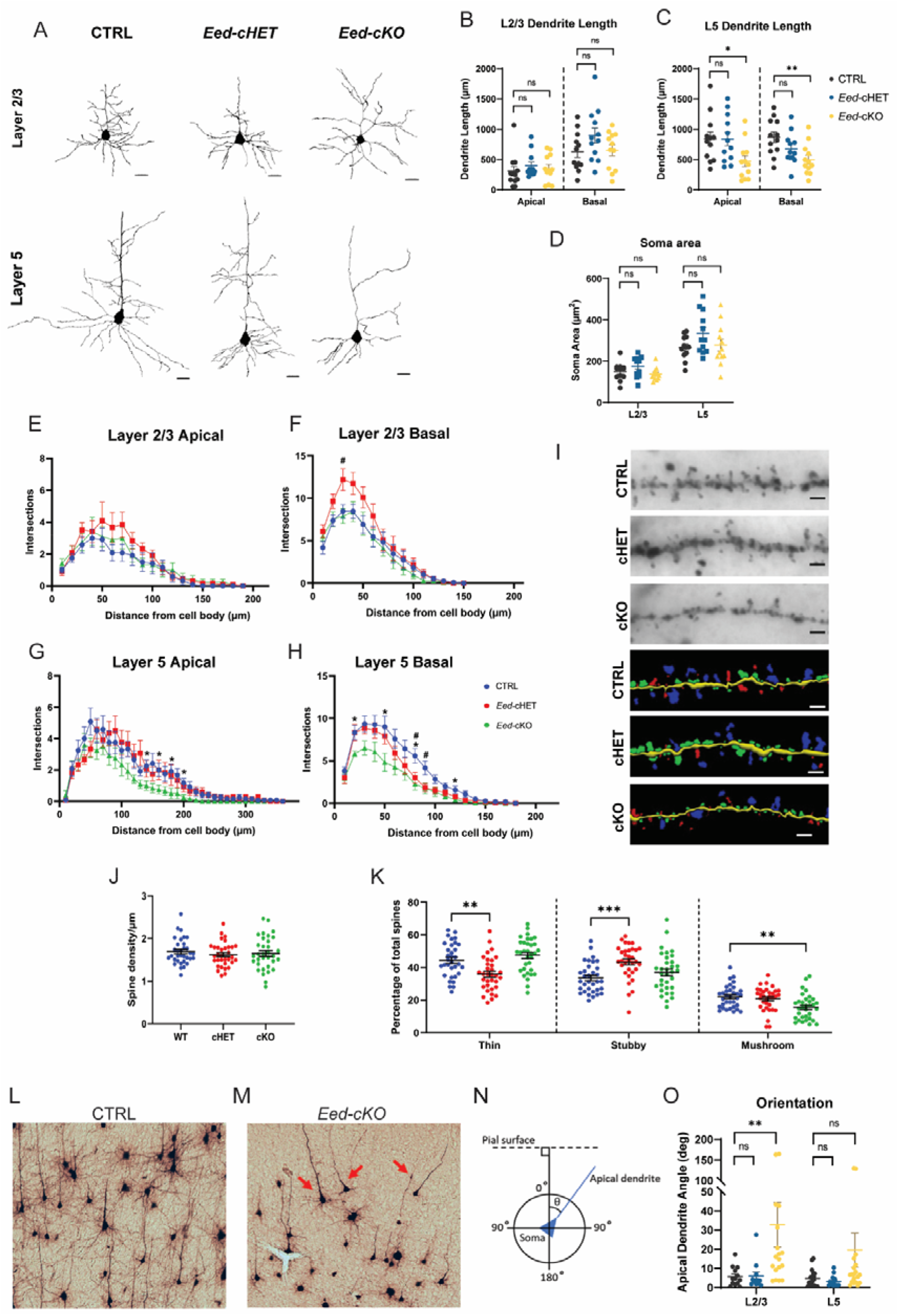
*Eed-cKO* pyramidal neurons have morphological deficits. Golgi-Cox analysis of CTRL, *Eed-cHet*, and *Eed-cKO* pyramidal neurons in layers 2/3 and 5 of the motor cortex. (A) Examples of pyramidal neuron morphology in layer 2/3 and 5 of the motor cortex in CTRL, *Eed-cHet*, and *Eed-cKO* mice. Scale bar = 20 µm. (B-C) Measurement of apical and basal dendrite length in layer 2/3 (B) and 5 (C) neurons. (D) Measurement of soma area in layer 2/3 and 5 pyramidal neurons. (E-H) Sholl analysis of dendrite complexity, which counts the number of dendrite branches in 10 µm intervals from the soma. # = p<0.05 (*Eed-cHet*), * = p<0.05 (*Eed-cKO*), two-way ANOVA (E-H). n = 12 neurons from 4 mice per genotype. (I-M) Analysis of spine morphology. (I) Examples of imaged (top) and digitally traced (bottom) dendrites with dendritic spines in CTRL, *Eed-cHet*, and *Eed-cKO* pyramidal neurons. Scale bar = 2 µm. (J) Density of dendritic spines was measured. (K) The percentage of thin, stubby, and mushroom shaped spines was also measured. ** = p<0.01, *** = p<0.001, **** = p<0.0001, one-way ANOVA with Dunnett’s multiple comparisons. n = 33 dendrites from 11 neurons per genotype across 4 CTRL, 4 *Eed-cHet* and 3 *Eed-cKO* mice. Each point in (J, M) represents an individual dendrite. (L-O) Analysis of apical dendrite orientation. (L, M) Examples of pyramidal neuron orientation in CTRL (L) and *Eed-cKO* (M) cortex. In the CTRL cortex, pyramidal neurons were orientated parallel to each other, however in the *Eed-cKO* cortex, there were several pyramidal neurons which were not parallel to each other (red arrows). (N) Schematic of how orientation was measured. The angle of the apical dendrite was measured relative to the pial surface. (O) Quantification of apical dendrite orientation. ** = p<0.01, ns = not significant, Kruskal-Wallis test with Dunn’s multiple comparisons. Layer 2/3: n = 12 CTRL, 12 *Eed-cHet*, and 17 *Eed-cKO* neurons from 4 mice per genotype. Layer 5: n = 24 CTRL, 15 *Eed-cHet*, and 23 *Eed-cKO* neurons from 4 mice per genotype.

During the Golgi-Cox analysis we observed that a small proportion of *Eed-cKO* pyramidal neurons were misoriented. For example, some neurons appeared to be inverted, with primary apical dendrites projecting towards the basal surface of the cortex instead of the pial surface (Figure 5O-P). To quantify the orientation of pyramidal neurons, the angle of the primary apical dendrite was measured relative to the pial surface (Figure 5N-P). There was no difference in orientation in *Eed-cHet* neurons compared to control, or in layer 5 *Eed-cKO* neurons. However, in layer 2/3 *Eed-cKO* neurons, the median apical dendrite angle was higher when compared to control neurons (Figure 5P). Hence, the apical dendrites of *Eed-cKO* neurons tended to be less perpendicular to the pial surface than in control neurons. This suggest that *Eed-cKO* pyramidal neurons might have deficits in apical dendrite guidance, causing abnormal neuron orientation. Overall, these results demonstrate that both *Eed-cHet* and *Eed-cKO* mice have deficits in dendritic arborization in pyramidal neurons, albeit of different nature. *Eed-cHet* neurons have a divergent phenotype to *Eed-cKO* neurons, a finding that is supported by the limited overlap in DEGs between *Eed-cHet* and *Eed-cKO* neurons.

### *Eed-cKO* mice have deficits in global white matter connectivity

Our findings of compromised neuronal structures (Figure 5) and forebrain commissures (Figure 1) suggested that cortical connectivity may also be aberrant in *Eed-cKO* mice. To investigate this, we utilized diffusion tensor magnetic resonance imaging (DTMRI) to reconstruct white matter tractography of the brain. First, we used DTMRI to examine metrics that provide information about the microstructural properties of white matter - fractional anisotropy (FA), axial diffusivity (AD), and radial diffusivity (RD)^41^. These metrics can be impacted by various changes in the tissue, such as alterations in tissue density, myelination, or composition of the extracellular matrix. Surprisingly, there was no significant difference in FA, RD, or AD in the corpus callosum, hippocampal commissure, or anterior commissure in *Eed-cHet* or *Eed-cKO* mice (Figure 6A-L). Voxel-based morphometry did detect changes in FA in several grey matter regions in the *Eed-cKO*, including the dentate gyrus and cortical plate (Supplementary 5). Nevertheless, the lack of changes in FA, RD, or AD in the commissures suggest that white matter integrity is not severely compromised in *Eed-cHet* or *Eed-cKO* mice. Interestingly however, we detected significant reductions in apparent fibre density^42^ in *Eed-cKO* mice, particularly in the internal capsule, hippocampal commissure, and in white matter tracts connecting the thalamus and globus pallidus (Figure 6M). By contrast, fibre cross section^42^ was not affected. Together, this suggests that they white matter tracts may be relatively normal, albeit less dense, in *Eed-cKO* mice.

**Figure 6.**
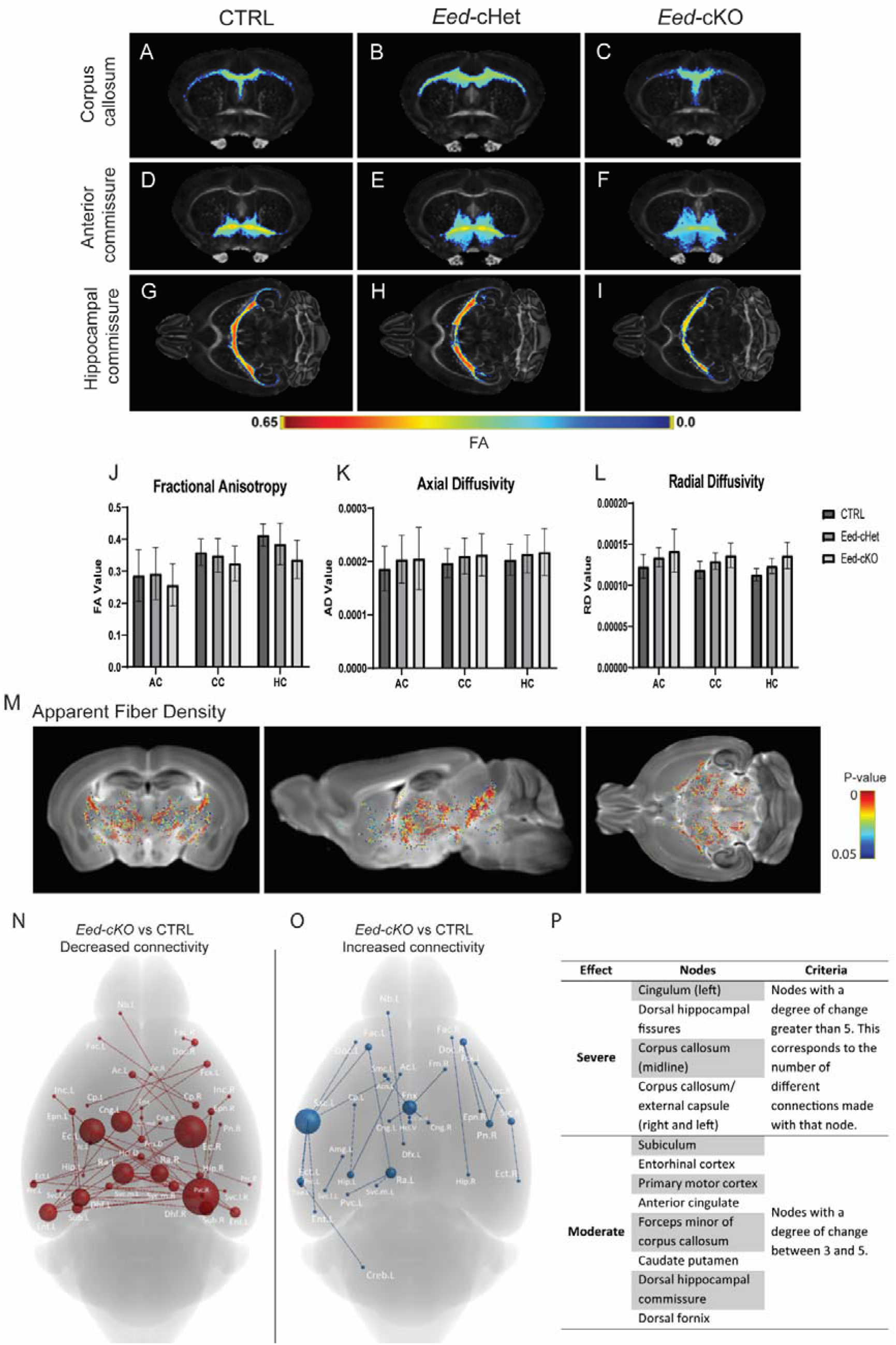
*Eed-cKO* mice have deficits in global white matter connectivity. (A-L) DTMRI metrics in the forebrain commissures of CTRL, *Eed-cHet*, and *Eed-cKO* mice. (A-R) Fractional anisotropy of the corpus callosum (A-C), anterior commissure (D-F), and hippocampal commissure (G-I) in CTRL, *Eed-cHet*, and *Eed-cKO* brains. There was no significant difference between genotypes. (J-L) Measurement of fractional anisotropy (J), axial diffusivity (K), and radial diffusivity (L) in the anterior commissure (AC), corpus callosum (CC), and hippocampal commissure (HC) There was no significant difference between genotypes, t-test. (M) Significant changes in apparent fibre density were observed in *Eed-cKO* mice compared to CTRL. (N-P) Connectome analysis of *Eed-cKO* brains. Graphic of key brain regions (“nodes”) which have decreased (N) or increased (O) connections to other regions in *Eed-cKO* mice. (P) A list of severely and moderately affected nodes in *Eed-cKO* mice. *Eed-cHet* brains are not displayed as there were no significant differences. n = 7 CTRL, 8 *Eed-cHet*, 5 *Eed-cKO*. See also Supplementary 5 for voxel-based morphometry analysis of FA.

How does this affect global connectivity in *Eed-cKO* mice? To investigate this we performed 3D reconstruction of the neural connections in the brain, allowing global connectivity to be investigated^43^. There was no significant change in the connectome detected between *Eed-cHet* and control mice. However, *Eed-cKO* mice had numerous regions with significantly altered connectivity compared to controls. Decreased connectivity was most common (Figure 6N), but there were also regions with increased connectivity (Figure 6O). The most severely affected regions included the cingulum, dorsal hippocampal fissures, and corpus callosum. The subiculum, entorhinal cortex, primary motor cortex, anterior cingulate cortex, caudate putamen, and dorsal fornix were also moderately affected (Figure 6P). We observed that many of the decreased connections involved interhemispheric connections, which was likely related to the smaller corpus callosum and hippocampal commissure.

### Loss of *Eed* results in mild behaviour alterations in the adult

Given the severe morphological phenotypes in *Eed-cKO* mice, as well as the distinct phenotypes within the *Eed-cHet*, we hypothesised that these mice would exhibit severe behavioural defects. To investigate this, we first performed the light-dark box test, which is used to investigate anxiety vs exploratory behaviours (Figure 7A-D)^44^. Interestingly, despite the changes in connectivity observed between limbic structures implicated in anxiety (Figure 6N-P), *Eed-cHet* and *Eed-cKO* mice did not spend more time in the light or dark component compared to controls (Figure 7B), although *Eed-cKO* mice did have reduced frequency in the number of times they moved between compartments (Figure 7C). Similarly, we did not observe any anxious behaviours in *Eed-cHet* or *Eed-cKO* mice in the open field test (Supplementary 6A-F)^45^. Anxiety/exploratory behaviours were further investigated with the elevated plus maze assay (Figure 7E-F)^46^. Surprisingly, *Eed-cKO* mice spent more time in the open arms than controls, suggesting that *Eed-cKO* mice may have less fear of open spaces than control mice, or that they are incapable of perceiving the danger (Figure 7F).

**Figure 7.**
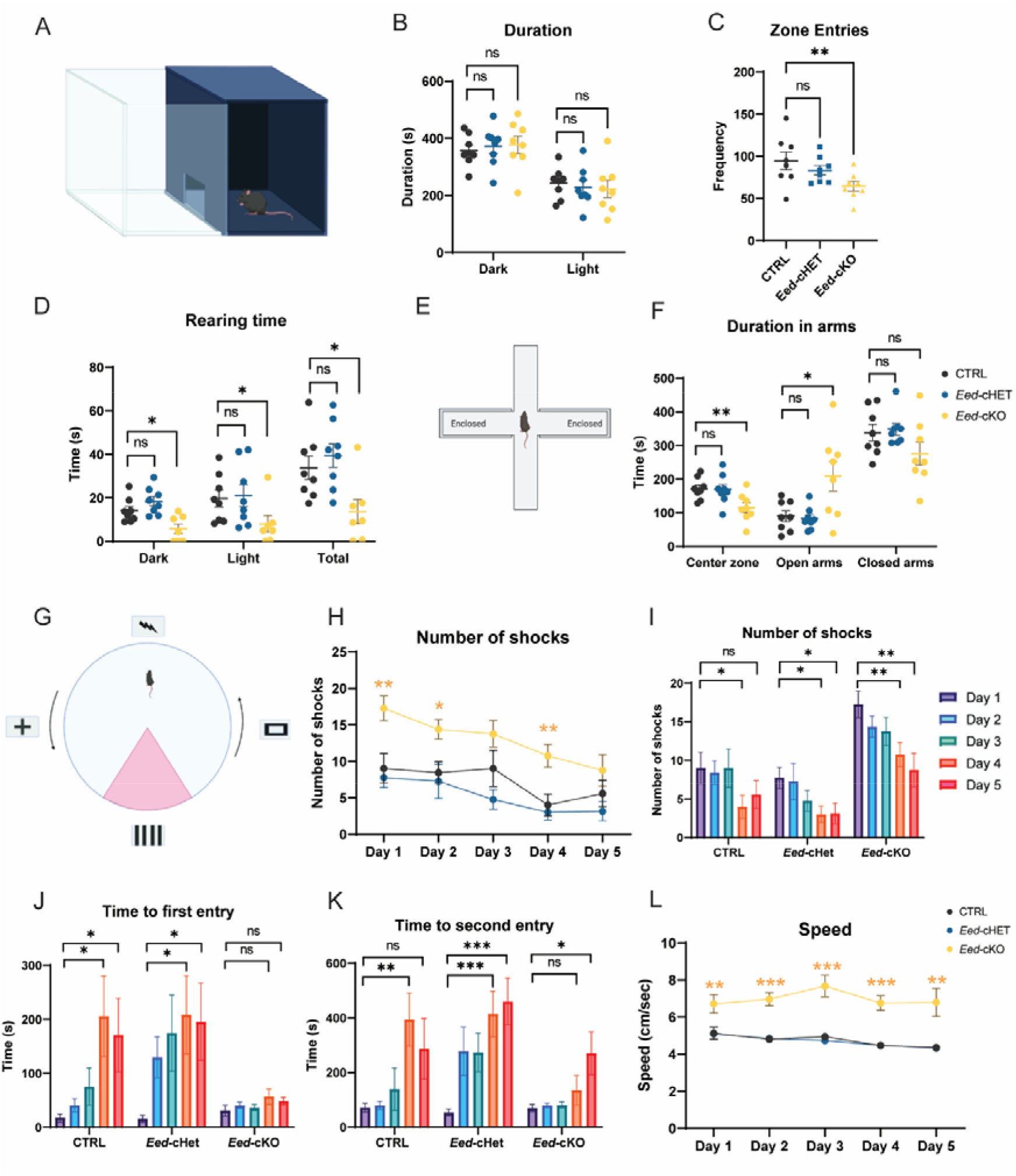
Loss of *Eed* results in mild behaviour alterations in the adult. (A-D) Light/dark box test. (A) Schematic of the light/dark box arena, which consists of a light component and a dark component separated by a small opening. (B-D) Duration in outer and inner zone (B), number of zone entries (C), and time spent rearing (D) was measured. There was no significant difference in time spent on other activities (ambulatory, stereotypic, jumping, resulting; data not shown). (E-F) Elevated plus maze test. (E) Schematic of the elevated plus maze arena, which consists of a plus (+) shaped arena suspended 1 m above the ground. There were two ‘enclosed’ arms with walls and two ‘open’ arms without walls. (F) Duration in centre zone, open arms, and closed arms was measured. There was no significant difference in frequency in arms, latency to arms, distance moved, average speed, duration not moving, or frequency of head dips (data not shown). (G-L) Active place avoidance test. (G) Schematic of the active place avoidance arena, which consists of a rotating arena with an invisible shock zone (red) in a fixed region, and fixed visual cues on the walls of the room. (H) Number of shocks is displayed in graphs for inter-genotype (H) and intra-genotype (I) comparison. Time to first (J) and second (K) shock and average speed (L) are also shown. In (H and L), yellow asterisks indicate statistical significance for *Eed-cKO* compared to CTRL, and blue asterisks indicates statistical significance for *Eed-cHet* compared to CTRL. *** = p<0.001, ** = p<0.01, * = p<0.05, one-way ANOVA. n = 8 CTRL, 8 *Eed-cHet*, 8 *Eed-cKO* (B, C, F) or 7 CTRL, 8 *Eed-cHet*, 8 *Eed-cKO* (H-L). See also Supplementary 6 for further details on the active place avoidance test, as well as additional behaviour tests.

Next, we assessed cognitive flexibility with the dynamic strategy-shifting test (DSST). Herein, mice are tasked to press a lever based on various visual and non-visual cues for a food reward. Despite the severe range of phenotypes at a cellular and molecular level in the *Eed-cKO* mice, there was no significant difference between genotypes, suggesting that cognitive flexibility was not compromised in *Eed-cHet* or *Eed-cKO* mice (Supplementary 7). *Eed-cHet* and *Eed-cKO* mice also do not display any deficits in sociability, as there were no significant differences in *Eed-cHet* or *Eed-cKO* mice in the three-chambered social interaction test (Supplementary 6G-M)^47^.

The severe hippocampal phenotype observed in cKO animals (Figures 1 and 2) led us to postulate that learning and memory may be abnormal in our cKO mice. To investigate this, we used an active place avoidance paradigm (Figure 7G-L, Supplementary 6K-O)^48^. The CTRL and *Eed-cHet* mice both received fewer shocks on later days of the trial, indicative of learning. To our surprise, although *Eed-cKO* animals received consistently more shocks each day over the 5-day protocol, they did exhibit learning, as evidenced by fewer shocks per day on day 5 compared to day 1 (Figure 7I). Closer analysis revealed that *Eed-cKO* mice did not exhibit a significant change in the time to the first shock each day – a proxy for long term memory – (Figure 7J). However, they did show improvements in short term memory, as measured by the time to the second entry to the shock zone (Figure 7K). The speed of the *Eed-cKO* animals was also increased over all 5 days of the trial. Again, these data point to an unexpected flexibility in our *Eed-cKO* mice; despite having severe hippocampal abnormalities, these mice were still able to learn in a hippocampal-dependent behavioural task. This is in stark contrast to other models that have similar hippocampal deficits and reduced cortical size, such as mice lacking *Usp9X*^49^.

## Discussion

Here we have characterized the adult effects of embryonic PRC2 loss-of-function in the dorsal telencephalon. As anticipated from previous studies, *Eed-cKO* mice have microcephaly, with significantly smaller and underdeveloped cortical structures including the cortical plate, corpus callosum, and hippocampus. Surprisingly, although the number of Ctip2+ and Satb2+Ctip2 neurons (denoting cortical layer 2/3 and 4-6 neurons respectively) were decreased, the number of Foxp2+ neurons (normally expressed by only by layer 6 neurons) was increased compared to controls. The cortical deficits are likely linked to decreased proliferation of NPCs and precocious neurogenesis, as shown previously^19–22^.

We also find a novel role for *Eed* in regulating neuron identity, as *Eed-cKO* glutamatergic neurons have highly dysregulated expression of neuron-specific genes and ectopic expression of interneuron, glial, and peripheral genes. Similar findings of GABA/Glut co-expression have been identified in the hypothalamus of *Eed-cKO* mice^29^, indicating that PRC2 may regulate neuron identity in different regions of the brain. Surprisingly, gene expression of cortical oligodendrocytes and astrocytes were not strongly affected, indicating that *Eed*, and consequently PRC2, does not play a major role in regulating gene expression in these cell types. Whether PRC2 function is similarly required for the specification of interneurons remains an open question; the use of different Cre drivers in future will be needed to investigate this question.

Gene ontology analysis indicated that numerous neuronal functions were altered in both *Eed-cHet* and *Eed-cKO* mice, such as synapse function, axonogenesis, and dendrite development. In line with this, Golgi-Cox identified deficits in both *Eed-cHet* and *Eed-cKO* mice in dendrite arborization and dendritic spine morphology, indicating that synaptic connections were abnormal. Interestingly, we also found that a small proportion of *Eed-cKO* pyramidal neurons were misoriented, which may indicate that *Eed* plays role in growth cone guidance. This notion is further supported by the connectome analysis, showing both decreased and increased connections between several brain regions. The connectome changes may underlie the effects upon some of the brain regions indirectly affected by *Emx1-iCre* deletion of *Eed*, such as the thalamus, which is highly interconnected to the neocortex and therefore may be smaller due to reduced reciprocal connections to the neocortex^50^.

The behavioural analysis revealed several interesting but unexpected results. Considering their cortical and hippocampal phenotypes, *Eed-cKO* mice performed surprisingly well at many tasks, including tasks involving exploration, social interaction, learning and memory, and cognitive flexibility. These tasks are cognitively complex and involve multiple cortical and limbic structures, many of which are severely diminished in *Eed-cKO* mice. Perhaps subcortical structures may be compensating for the diminished cortical structures in *Eed-cKO* mice. Functional MRI may provide further insight into this. Alternatively, even though *Eed-cKO* cortical and hippocampal structures are diminished, they are still able to function well enough for these behaviour tests. Alternatively, neuronal activity in different brain regions could be investigated with immunostaining for early expressed genes such as *c-fos*^51^.

In *Eed-cHet* mice, we did not observe any major alterations in the structure, cytoarchitecture, or connectivity in the cortex, or in behaviour. However, *Eed-cHet* mice did have alterations in glutamatergic neuron gene expression and pyramidal neuron morphology. As such, it appears that while the phenotype of *Eed-cHet* mice is subtle, loss-of-function of one allele of *Eed* is enough to cause deficits in cortical development. Remarkably, *Eed-cHet* mice often had a divergent phenotype to *Eed-cKO* mice. For instance, only a very small proportion of DEGs overlapped between *Eed-cHet* and *Eed-cKO* neurons. Moreover, *Eed-cHet* pyramidal neurons had increased dendritic arborization in upper layers, which *Eed-cKO* pyramidal neurons did not show. *Eed-cHet* dendrites also had a higher proportion of stubby spines whereas *Eed-cKO* dendrites had a lower proportion of mushroom spines. Collectively, this suggests that that loss of one allele (*Eed-cHet*) vs two alleles (*Eed-cKO*) of Eed may have different epigenetic and gene expression consequences, pointing to a dose-sensitivity of the PRC2 complex. The heterozygous phenotypes observed in our *Eed-cHet* animals may have bearing upon the role of PRC2 in humans, where the three key PRC2 genes *EED*, *EZH2* and *SUZ12* are all haploinsufficient^52,53^. Moreover, the human Weaver, Weaver-like, Cohen-Gibson, and Overgrowth and Intellectual Disability syndromes are linked to heterozygous, possibly neomorphic or hypermorphic mutations in *EED*, *EZH2* or *SUZ12*^11–13^. These syndromes present with neurological defects, which involve delayed speech and psychomotor development, and intellectual disability. MRI analysis has not been widely conducted on these patients, but in the few cases where it has been used it has shown anatomical defects, including polymicrogyria and white matter volume loss^54,55^. A scenario is thus emerging where half-dose, no-dose and neo/hyper-morphic genetic defects in PRC2 genes all negatively affect mammalian brain development.

## Methods

### Mouse Model

All animals were used with approval from the University of Queensland Animal Ethics Committee (AEC approval number 165/19/ARC, 2022/AE000397, and QBI/191/20). All work was carried out in accordance with the Australian Code of Practice for the Care and Use of Animals for Scientific Purposes and the University of Queensland’s Institutional Biosafety Committee.

*Eed*^fl/fl^ mice were crossed with *Emx1*-iCre^+/-^ mice. This produced pups which were *Eed*^wt/fl^; *Emx1*-iCre^+/-^. Then, *Eed*^wt/fl^; *Emx1*-iCre^+/-^ mice were crossed with *Eed*^fl/fl^ mice. This produced pups which were *Eed*^fl/fl^; *Emx1*-iCre^+/-^ (*Eed-cKO*, homozygous deletion of *Eed*) and *Eed*^wt/fl^; *Emx1*-iCre^+/-^ (*Eed-cHet*, heterozygous deletion of *Eed*). This cross also produced pups which were *Eed*^fl/fl^; *Emx1*-iCre^-/-^ and *Eed*^wt/fl^; *Emx1*-iCre^-/-^, which did not have deletion of *Eed* and were used as littermate controls (CTRL). In all crosses, there were no difficulties with breeding and the genotypes of pups were expected the Mendelian ratios.

### Genotyping

Genotyping was completed for all associated mouse lines. DNA was extracted from tissue by submerging in lysis buffer (25mM NaOH, 0.2mM EDTA) for 30 minutes at 95°C, then neutralising with neutralisation buffer (40mM Tris-HCl). Genotyping was performed by running a PCR with Platinum Green Hot Start PCR Master Mix (ThermoFisher). Details on the PCR cycle conditions are including in Table 1 and details on primers are included in Table 2. The PCR products were run on a 2% agarose gel containing 0.5 μg/mL ethidium bromide at 100V for 20 minutes in TAE buffer. Gels were imaged using the AlphaImager Gel Imaging System (Alpha Innotech). For *Eed* genotyping, bands were 313 base pairs in size for the Wt allele, and 360 base pairs in size for the floxed allele. For *Emx1* genotyping, bands were approximately 600 base pairs for *Emx1*^+^ samples or absent on the gel for *Emx1*^-^ samples.

**Table 2:**
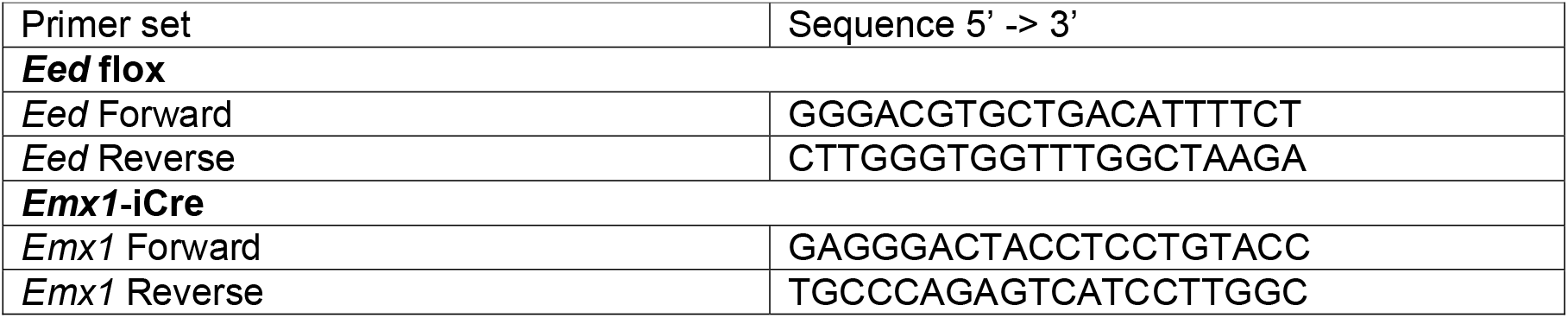
Genotyping primers.

### Perfusion

Adult mice were euthanized by intraperitoneal injection of 0.8 mL of 1:50 Lethabarb (Virbac). Incisions were made though the integument beneath the rib cage, the diaphragm, and then along the rib cage parallel to the sternum. The rib cage was folded back to expose the heart. A 27-gauge needle attached to a peristaltic perfusion pump was inserted into the left ventricle of the heart. The right atrium was cut to allow the flow of liquid. Approximately 15-20 mL of 1X PBS solution was pumped through the body until the liver was clear. Following this, approximately 30-40 mL of 4% paraformaldehyde (PFA) was pumped through. Once completed, the mouse was decapitated, then the brain was collected by making a longitudinal midline incision to the skull, fragmenting it into pieces which were removed. The brain was post-fixed in 4% PFA for 48 hours before storing in PBS at 4°C.

### Vibratome Sectioning

Fixed brains were embedded in 3% noble agar gel and sectioned with the Vibratome VT1000S (Leica) at 50 μm thickness as free-floating sections. Sections were stored in PBS with 0.02% sodium azide at 4°C.

### Haematoxylin

Vibratome-cut sections (50 µm thick) were mounted on SuperFrost Plus microscope slides (Menzel-Glasser, Thermo-Fisher Scientific) and dried at 37°C for approximately 5 minutes. Sections were then rehydrated in H_2_O for 1 minute, submerged in hematoxylin stain for 2.5 minutes, then gently rinsed with H_2_O for 1 minute.

The sections were dehydrated with the following steps: H_2_O for 1 minute, 70% ethanol for 2 minutes, 95% ethanol for 2 minutes, and 3x 100% ethanol for 2 minutes. The sections were then placed in xylene for 3 minutes, twice. Finally, the slides were mounted with DPX mounting medium. Imaging was performed using the Aperio SlideScope XT. Images were analysed with the Aperio ImageScope software.

### Immunofluorescent Staining

Vibratome-cut sections (50 µm thick) were mounted and dried on SuperFrost Plus slides and dried at 37°C for approximately 15 minutes. Antigen retrieval was performed with the Decloaking Chamber NxGen (Biocare Medical). The decloaking chamber was set up with 500 mL of mH_2_0 in the base tray, 230 mL of mH_2_0 in the side containers, and 230 mL of citrate buffer (made with 2.94 g sodium citrate in 1 L mH_2_0, pH 6 with HCl). Antigen retrieval was run at 95°C for 15 minutes. Once cooled, excess liquid was wiped off and the samples were outlined with PAP pen. Slides were washed 3x with PBS for 10 minutes.

Sections were incubated in blocking serum (0.2% animal serum and 0.02% Bio-Rad TritonX in PBS) for 60 minutes at room temperature. Then, the sections were washed 3x with PBS for 10 minutes. The primary antibody was diluted into serum block (Table 1), and sections were incubated in the primary antibody solution overnight at 4°C. The following day, sections were washed 3x with PBS for 5 minutes. Next, the secondary antibody was diluted in serum block (Table 1). Sections were incubated in secondary antibody for 60 minutes at room temperature, in the dark. Sections were washed 3x with PBS for 5 minutes. Then, sections were incubated in DAPI solution (1/1000 concentration) for 10 minutes at room temperature. Sections were washed 3x with PBS for 10 minutes. Coverslips were mounted with DAKO Fluorescent mounting medium and sealed with nail polish.

Slides were imaged with the Diskovery spinning disc confocal microscope or the Aperio Scanscope XT slidescanner and analysed with the FIJI software.

### Golgi Cox Staining

#### Preparation of Golgi-Cox solution

800 mL of Golgi Cox solution was prepared by combining three solutions, 5% Potassium dichromate (Solution A), 5% Mercuric chloride (Solution B), and 5% Potassium chromate (Solution C).

Solution A was prepared by dissolving 5 g of potassium dichromate in 100 mL of distilled H2O. Solution B was prepared by dissolving 5 g of mercuric chloride in 100 mL of distilled H2O, using a magnetic stirrer and a hot plate to dissolve the crystals. Solution C was prepared by dissolving 4 g of potassium chromate in 80 mL of distilled H2O. Solution C was further diluted by adding 200 mL of distilled H_2_O. Solution A was slowly mixed with Solution B, then added slowly into Solution C while stirring on a hot plate (low-medium heat). The solution was stirred until it was red in colour and cloudy. The solution was stored in the dark for 5 days, then filtered, to create the working solution, which is stable for approximately 14 days.

#### Histochemical staining

Perfusions were performed with 0.04% PFA. Freshly perfused brains were placed in approximately 30 mL of Golgi-Cox solution in a sealable specimen container. Brains were incubated in Golgi-Cox solution for 10 days at room temperature. After the first 5 days, the Golgi-Cox solution was replaced with fresh solution. After 10 days, the brains were placed in 30% sucrose and incubated for 4 days at 4°C. After 4 days, Brains were sectioned at a thickness of 150 µm using a vibratome. Sections were placed in 6 well plates containing 30% sucrose solution. The sections were then mounted onto SuperFrost Plus slides immersed in 1.25% gelatin and 0.1% chromium. Mounted tissue was dried for 2 days at 4 degrees Celsius in the dark.

#### Post-processing

Slides were rehydrated in H_2_0 for 2 minutes, then placed in 0.1M ammonia solution for 30 minutes in the dark. Then, the slides were rinsed twice with H_2_0 for 5 minutes each. The slides were placed in Fujifilm Automatic X-Ray fixer solution for 30 minutes in the dark. The slides were rinsed twice with H_2_0 for 5 minutes each, then dehydrated through series of ethanol solutions: 70%, 90%, 95%, 100%, 100% and 100% ethanol for 5 minutes each. The slides were placed in CXA solution (1:1:1 chloroform: xylene: alcohol) for 10 minutes, then placed in xylene two times, 5 minutes each. Finally, the slides were mounted with Dibutylphthalate Polystyrene Xylene (DPX) and coverslips were added.

#### Imaging and Analysis

Imaging was performed with the Nikon Upright Stereology microscope. Layer 2/3 and Layer 5 pyramidal neurons from the motor and somatosensory cortices were imaged. Pyramidal neurons were identified by their typical triangular shaped soma and distinct apical dendrites which project towards the outer surface of the cortex. To be selected, neurons had to be clearly stained, and not visibly damaged. Neurons too close to other stained neurons that their individual structures could not be defined were not selected. Single neurons were imaged as Z-stacks with a step size of 2 μm at 40X magnification. Z-projections were then produced from the stacks for tracing.

The Neurolucida 360 software (version 2019) was used to digitally trace the cell soma and dendrites of the imaged neurons^56^. Dendrites extending from the apex of the soma were classified as apical and all other dendrites were classified as basal. Traces were then imported into Neurolucida Explorer software (version 2019), where data on total apical and basal dendrite length and cell soma area were generated. The software was also used to perform Sholl analysis on the apical and basal dendrites of the traces. Concentric rings centred on the cell soma and spaced at 10 μm intervals were overlaid onto the traces and the number of intersections automatically counted.

#### Apical dendrite orientation analysis

For the orientation analysis, the whole motor and somatosensory cortices of each brain were imaged as Z-stacks with a step size of 5 μm at 10X magnification. Images stacks were then loaded into FIJI Image J where a 0.1 mm2 area grid was overlaid onto them. All suitable pyramidal neurons within a randomly selected square in Layer 2/3 and Layer 5 were selected for analysis. If a square had less than 3 suitable neurons in it, another square would be selected until at least 3 neurons had been analysed. Pyramidal neurons were defined as suitable according to the same criteria as the neuron tracing. Apical dendrite orientation was quantified in FIJI Image J as described^57^. A line was drawn extending out from the centre of the cell soma that is perpendicular to the outer surface of the cortex. Another line was drawn from the centre of the soma along the direction of the apical dendrite. The angle between these two lines was then measured.

#### Dendritic spine quantification

To analyse dendritic spines, terminal basal dendrite branches of Layer 2/3 pyramidal neurons from the motor cortex were imaged as Z-stacks with a step size of 0.3 μm using an 100x oil objective lens. Neurons and branches were selected at random. Pyramidal neurons were identified according to the same criteria as the neuron tracing. Terminal branches had to be at least secondary order, at least 10 μm in length and at least 10 μm from the cell soma to be used for analysis. Z-projections were produced from the stacks for tracing. Using Neurolucida 360 software (version 2019), selected branches were digitally traced. The “detect spines” feature was then used to automatically detect all dendritic spines associated with the branches (Detector settings: outer range = 2.5 μm, minimum height = 0.3 μm, detector sensitivity = 110%, minimum count = 10 voxels). The “classify spines” feature was then used to automatically classify all spines as thin, stubby, or mushroom spines. Spine types were defined as follows: thin spine = head-to-neck diameter ratio <1.1, spine length-to-head diameter ratio >2.5, and filipodium length <3 μm OR head-to-neck diameter ratio >1.1, mushroom head diameter < 0.35 μm, and filipodium length <3 μm, stubby spine = head-to-neck diameter ratio <1.1 and spine length-to-head diameter ratio <2.5, mushroom spine = head-to-neck diameter ratio >1.1, mushroom head diameter >0.35 μm. Traces were then imported into Neurolucida Explorer software (version 2019) where data on spine density, length and head diameter were generated.

### DTMRI

#### Tissue preparation

For DTMRI analysis, a total of 24 adult mice (7 CTRL, 8 *Eed-cHet* and 5 *Eed-cKO*) were used. Mice were perfused as described above; however image data was acquired while the brain was still inside the skull to perverse native spatial relationship and to make sure all brain tissue was intact. Samples were further post-fixed in 4% PFA for 24 hours, and then stored in 0.1 M PBS with 0.02% sodium azide (Sigma-Aldrich, USA). Before MRI scanning, the skulls were immersed in 0.1 M PBS with 0.2% gadobutrol (Gadovist, Bayer, Ontario, Canada) for 8 weeks to enhance MRI contrast.

#### Diffusion MRI Acquisition

Brain data was acquired using the Bruker Ultrashield Plus 700 WB Avance NMR spectrometer (16.4 T vertical bore animal microimaging system). It was equipped with a 15 mm linear surface acoustic wave coil, a micro2.5 gradient insert that provides a 2.5 mT/m/A gradient strength, and a 60 A gradient amplifier. Two sets of scans were obtained: a high-resolution 3D T1 weighted Fast Low Angle Shot (FLASH) for structural information imaging; and 3D Stejskal-Tanner DWI spin-echo in multiple directions for fibre tracking purposes. Samples were scanned using a high resolution FLASH T1 weighted sequence and a DWI protocol as illustrated in Table 4 and Table 5 below.

**Table 3:**
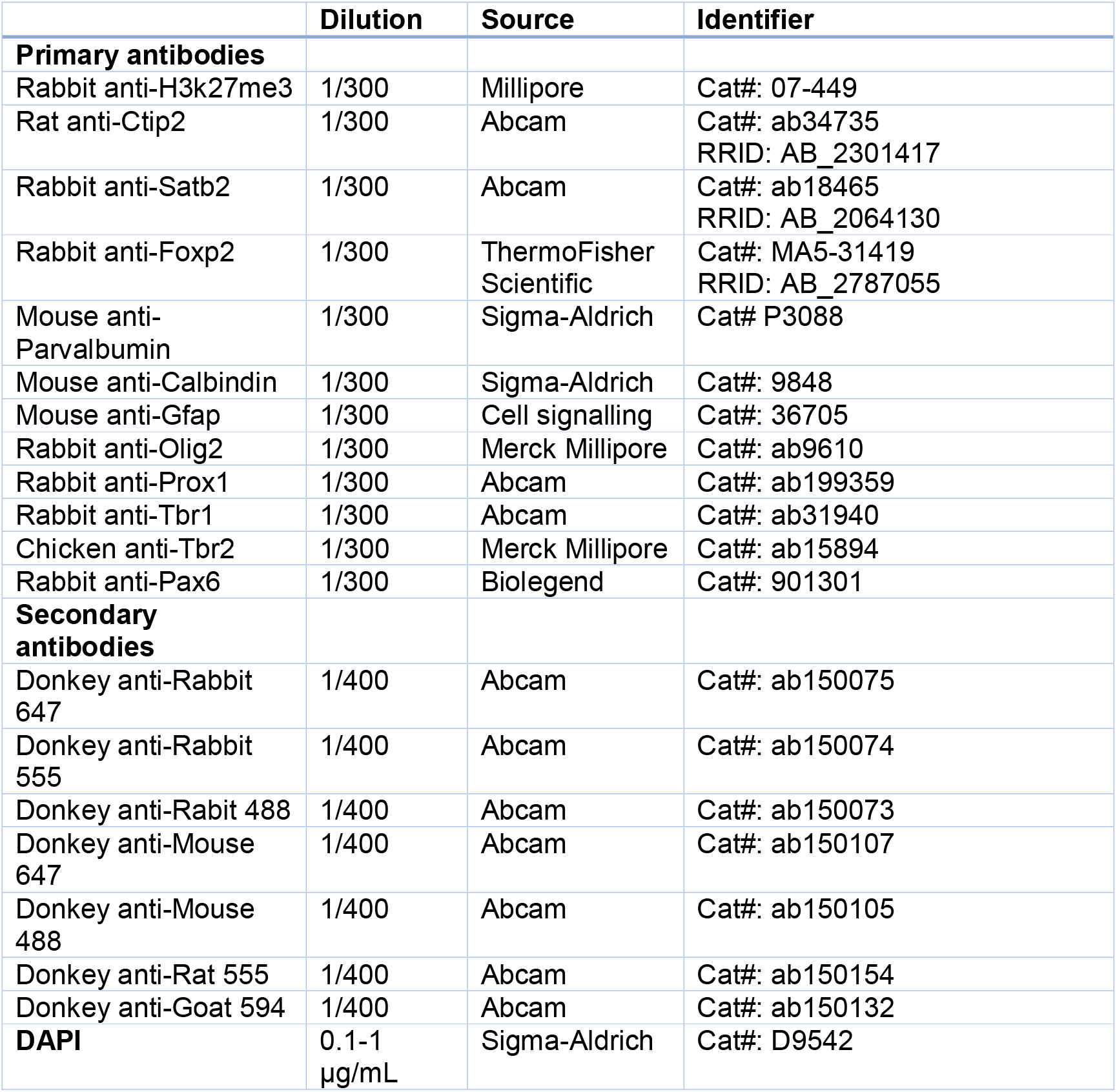
List of antibodies.

**Table 4:** High resolution T1 imaging sequence parameters.

| High resolution T1 |  |  |  |  |  |
| --- | --- | --- | --- | --- | --- |
| TR | TE | Matrix Size | FOV | Resolution | Bandwidth |
| 50 ms | 12 ms | 654x433x333 | 19.6x13x10 mm | 30 $\mu\text{m}$ Isotropic | 50 KHz |

**Table 5:** DWI sequence parameters.

| DWI |
| --- |

| TR | TE | *b* value | Matrix Size | FOV | Resolution | Bandwidth |
| --- | --- | --- | --- | --- | --- | --- |
| **200 ms** | 21 ms | 5000 s/mm<sup>2</sup><br>3x*b*<sub>0</sub> 30<br>directions | 196x130x100 | 19.6x13x10 mm | 100 µm | 50 KHz |

#### Data pre-processing and analysis

DWI data were zero-filled by a factor of 1.5 prior to Fourier transform. Non-brain tissue was removed through masking brain tissue only using ITKsnap software package^58^ then the diffusion data were denoised and bias-field corrected using both MRtrix and ANT’s software packages^59,60^. MRtrix was used to calculate the response function as well as the iFOD2 map^60^.

A 106 regions of interest (ROIs) parcellation images created from an MRI based atlas of adult C57BL/6 mouse from both the Center for Advanced Imaging at the University of Queensland and Susumu Mori’s lab of Johns Hopkins University were used for probabilistic tractography^61^. Figure 1 illustrates how the data was processed to generate the structural connectome. Both the atlas and the parcellation images were registered to each subject’s b_0_ image using FMRIB Software Library (FSL) Linear Image Registration Tools (FLIRT) rigid-body registration, followed by affine and nonlinear deformation using ANTs^62–64^. The ROIs were defined as nodes between which the edges were computed.

Both the template and atlas were transformed into the subject space using the subject b=0 image. Subsequently whole brain probabilistic tractography was performed and processed to generate the structural connectome.

#### Structural connectome and VBM analysis

The connectome was studied network-wise by graph theory analysis and connection-wise using the Network-Based Statistic Toolbox (NBS)^43,65^. For NBS, connectomes were compared over multi-thresholds ranging between t = 2.1 to t = 6.5 with an increment of 0.1 each time to compute the FWER-corrected p-value for each component. For graph theory analysis, network metrics were measured over 10 thresholds ranging between t = 0.05 to t = 0.5 of the network sparsity. These metrics include degree centrality, betweenness, small worldness, and modularity.

#### Volumetric analysis

The volumetric change in *Eed-cHet* and *Eed-cKO* mice compared to CTRL mice was also investigated in 20 regions using a comprehensive three-dimensional digital atlas by Ma et al^66^. These regions are shown in **Error! Reference source not found.**

#### DTMRI Metrics

DTMRI maps were calculated using MRTrix3, and DTMRI metrics (FA, AD, and RD) were measured using the FSL software. To investigate the white mater integrity, DTMRI metrics were measured from three main regions of interests, these were the corpus callosum, hippocampal commissure, and anterior commissure. Fixel-based analysis was performed using MRTrix3 to investigate WM fibre density and morphology^42^. FOD maps were used to create an unbiased FOD 4D template then each subject’s FOD map was registered to the template and warped to the template’s space. These warped-FOD images were then segmented to estimate fixels and their AFD while “warp” images were used to compute the fibre cross-section. Probabilistic Tractography was performed on the FOD template to generate a whole-brain tractogram. This tractogram was used to generate a fixel-fixel connectivity matrix. Finally, fixel data was smoothed then statistical analysis was performed using connectivity based fixel enhancement for each measure (AFD and cross-section)^42^.

Voxel-based morphometry (VBM) analysis was performed using Threshold-Free Cluster Enhancement (TFCE) with controlled family-wise error (FWE) rate to detect structures of the brain that were most significantly affected by *Eed* ablation^67^. Multivariate templates were created using ANTs^62–64^. DTMRI-derived maps were registered together in a non-biased space firstly using linear transformation (rigid and affine) followed by a deformable transformation with higher degrees of freedom^68^. FSL randomise was then performed to calculate the FWE-TFCE test statistic maps and a group of p value maps^67^.

### High Resolution Episcopic Microscopy (HREM)

#### Tissue collection and processing

Brains were extracted from the skull and fixed in Bouin’s fixative for 24 hours. Fixed brains were washed in PBS and then stored in PBS at 4°C. Brains were then dehydrated by submerging them in a series of methanol solutions: 10%, 20%, 30%, 40%, 50%, 60%, 70%, 80%, 90%, and 95% for 2 hours each, then 100% three times for 1 hour, 1 hour, and 2 hours, respectively. Brains were then infiltrated with JB-4 Plus methacrylate resin dye containing eosin B and acridine orange for 13 days at 4°C in a rotary tube mixer, changing to fresh resin dye mix on the fifth day. Brains were embedded individually in moulds using fresh JB-4 Plus resin dye mix. Resin blocks were allowed to polymerise and were incubated at 90°C for 5 days, or until hardened. The resin-embedded brains were mounted onto the HREM machine to be sectioned and imaged. The brains were serially sectioned at a thickness of 5 μm and imaged using a 470 nm LED light. Images of the block face were captured following every section, using a fluorescent 470 channel.

#### Whole brain volumetric analysis

Raw image stacks of coronal sections of whole mouse brains were generated by high resolution episcopic microscopy. For each brain, every fourth image was selected and then compiled into three-dimensional whole brain reconstructions using Amira software (version 2021.1). The software’s ‘segmentation editor’ feature was then used to segment the reconstructions into several different brain structures. These were the neocortex, olfactory areas of the cortex (includes the anterior olfactory nucleus, taenia tecta, dorsal peduncular area, piriform area, nucleus of the lateral olfactory tract, cortical amygdala area, piriform-amygdala area and the postpiriform transition area), olfactory bulb (includes the main and accessory bulb), entorhinal cortex, hippocampus, corpus callosum, anterior commissure (excluding the olfactory bulb portion), amygdala (includes the lateral, basolateral and basomedial amygdala), caudoputamen, globus pallidus, lateral ventricles, thalamus, and hypothalamus. The Allen Mouse Brain Reference Atlas was used as a reference to segment the structures. Amira software was then used to generate volumetric data for each structure. Note that for the olfactory bulb, only the data for a single olfactory bulb was used for analysis. This was because not all brains measured had both olfactory bulbs intact. For brains that did have both intact, data from the largest bulb was used. 3 mice per genotype were used for volumetric analysis.

### Single nuclei-RNA-Seq

Homogenisation buffer (HB) and wash media (WM) were prepared and placed on ice. The homogenisation buffer was composed of 250 mM sucrose, 25 mM KCl, 5 mM MgCl2, 10 mM Tris buffer (pH 8), 1 µM DTT, 1x Protease inhibitor, 0.1% (v/v) Triton X-100 10%, and 40 U/mL RNaseIn, diluted in nuclease-free water. The wash media was composed of 1% BSA and 100 U/mL RNasIn, diluted in phosphate buffered saline (PBS). The Dounce homogenizer was precooled on ice.

Tissue was dissected and immediately transferred into the Dounce homogenizer, filled with 1ml of HB. The tissue was homogenised with 10 strokes of the loose pestle, followed by 15 strokes of the tight pestle. The homogenate was transferred to a prechilled 15 ml tube and 4ml of cold HB was added, then incubated for 5 minutes on ice. The homogenate was then strained through a 40µm cell strainer prewetted with 0.5ml HB, into a 50ml tube. The strainer was washed with 0.5ml HB. The homogenate was centrifuged for 5 minutes at 500g in a swing bucket. The cell pellet was resuspended in 2ml WM, and DAPI was added at 1:5000 concentration in 1ml WM.

The cell suspension was submitted for FACS sorting, using DAPI to filter nuclei from debris. Finally, the samples were submitted for sequencing with the 10x Genomics Chromium platform. The raw data was remapped from a whole cell genome to a pre-mRNA genome using Cell Ranger. Analysis was performed with the Seurat toolkit for single cell genomics on R Studio^69^. Firstly, CTRL, *Eed-cHet*, and *Eed-cKO* data was merged into one Seurat object. Cells with <2% mitochondrial RNA were filtered out as these were likely to be low-quality cells such as dead or dying cells. The data was normalised with both SCTransform^70^ (for cluster calculations) and NormaliseData (for data visualisation). Cells were then clustered with the standard Seurat workflow (RunPCA, RunUMAP, FindNeighbors, FindClusters) (dims = 1:30, resolution = 0.2). Clusters were identified based on canonical gene expression, and by using Linnarsson lab scRNA-seq data of the mouse cortex and hippocampus for reference of gene expression in different cell types^71^. Clusters with unusual nFeature, nCount, or abnormal gene expression were filtered out as these were likely to be doublets or abnormal cells. Differentially expressed gene (DEG) analysis performed with FindMarkers (wilcox test, logfc.threshold = 0.2). Then, glutamatergic neuron clusters were isolated by subsetting them and forming new Seurat object. The workflow for analysing glutamatergic neurons was like that described above. Volcano plots were generated with the EnhancedVolcano package^72^. Finally, Gene ontology (GO) analysis was performed with the clusterProlifer function enrichGO^73^.

### Behaviour

All mice were group-housed (2-5 mice per cage) within individually ventilated cages under a 12-h light/dark cycle (lights on 0700h) in a room maintained at constant temperature (21 degrees) and humidity (60%) within a PC2 animal facility.

Three different cohorts of mice underwent behaviour analysis. The first and second cohort both consisted of 4 females and 4 males per genotype, amounting to a total of 8 CTRL, 8 *Eed-cHet*, and 8 *Eed-cKO* mice. The third cohort consisted of 4 females and 5 males per genotype, amounting to a total of 9 CTRL, 9 *Eed-cHet*, and 9 *Eed-cKO* mice. Throughout the testing, males and females were kept separate and the arena was thoroughly wiped with 70% ethanol between testing each mouse to ensure that no smells would affect the behaviour of the next mouse.

The first cohort underwent SHIRPA and the 3-chambered social interaction test. The second cohort underwent the activity monitor, light/dark box, elevated plus maze, and active place avoidance tests. Finally, the third cohort underwent the cognitive flexibility test. The order of tests for each cohort was designed to begin with the least stressful tests, to minimize the impact of previous experiences on the latter tests.

#### SHIRPA

Mice were screened for gross neurological deficits using SHIRPA (Smith Kline Beecham Pharmaceuticals; Harwell, MRC Mouse Genome Centre and Mammalian Genetics Unit Imperial College School of Medicine at St. Mary’s; Royal London Hospital, St. Bartholomew’s and the Royal London School of Medicine phenotype assessment) screen^74^. Mice underwent a series of tests and observations to assess their health, sensory and motor function, neurological reflexes, and autonomic functions ^74^. No neurological deficits were found in CTRL, *Eed-cHet*, or *Eed-cKO* mice.

#### Activity Monitor

The activity monitor consists of an enclosed arena in which mouse movements were tracked with infrared sensors (**Error! Reference source not found.** A, B). Factors such as ambulatory movement (walking or running), stereotypic movement (small movements such as scratching), rearing, jumping, and resting were recorded. Additionally, the location of the mouse in the arena was recorded. Each mouse was recorded in the activity monitor arena for 30 minutes. The lighting at the level of the arena was 80 lux.

#### Light/Dark Box

The light/dark box consists of two compartments separated by a small door. The ‘light’ compartment was brightly lit with an open roof, and the ‘dark’ compartment was covered and dark (**Error! Reference source not found.**A). Like the activity monitor, mouse activity was recorded with infrared sensors. Ambulatory, stereotypic, resting, rearing, and jumping movements were recorded. This test was performed for 10 minutes. The lighting in the light compartment of the arena was 80 lux.

#### Elevated Plus Maze

The elevated plus maze arena is in the shape of a plus symbol (+), with two open and two enclosed arms (**Error! Reference source not found.**A). The arena is placed on a pedestal 1m above the ground. Normally mice are averse to open spaces and prefer the enclosed arms. Hence, this test investigates anxious vs exploratory behaviours. This test was performed for 10 minutes per mouse. The lighting of the arena was 80 lux.

#### 3-Chambered Social Interaction Test

The three-chambered apparatus consisted of three chambers, each compartment was 20×30×30cm, with a wire frame steel cage in the left and right chambers (**Error! Reference source not found.**A). Firstly, the subject mouse was placed in the centre of the apparatus and allowed to habituate for 5 minutes. After 5 minutes, the subject mouse was confined to the centre chamber using dividers. A conspecific mouse of the same sex was placed in either the left or right chamber within a wire framed steel cage. The dividers were removed, and the subject mouse was allowed to move freely between the three chambers for 5 minutes. Next the subject mouse was confined to the central chamber again, and a second conspecific mouse of the same sex was placed in the remaining wired framed steel cage. The dividers were removed, and the subject mouse was able to move freely between the three chambers for a further 5 minutes. Video recordings were coded and analysed blind to treatment using Ethovision XT (Noldus Information Technology, NLD) to track the centre of mass (body). The cumulative duration and frequency of the mouse to enter each zone was measured as well as the amount of time spent near the conspecific mouse. The lighting at the level of the three-chambered apparatus was 100 lux.

#### Active Place Avoidance

The APA (manufactured by Bio-Signal Group) task was employed to assess hippocampal-dependent spatial learning^75^. The APA apparatus consists of an elevated spherical arena (77cm in diameter and 32cm in height) with a grid floor (**Error! Reference source not found.**A). The arena rotated clockwise at 1rpm. The task of the mouse was to avoid an invisible shock zone by using four external cues which were hung on nearby walls of the room. If the mouse entered the shock zone, it would receive a mild foot shock (500 ms, 60 Hz, 0.5 mA) via the grid floor. The mouse received a further shock if it did not escape the shock zone [same intensity with 1.5-second intervals]. An overhead camera was used to track the mouse and a computer-based software was used to analyse the image and to deliver the mild shock appropriately (Tracker, Bio-Signal Group Corp., Brooklyn, NY). The experiment was conducted over 6 consecutive days, habituation (Day 0, without shock for 5) and training (Day-1 to 5, with shock 10 minutes/day). Behavioural parameters such as “number of shocks received”, “latency to enter into the shock zone”, “max time avoid”, and “distance travelled” were analysed at the end of the experiment using software (Track Analysis) provided by the Bio-Signal Group. The lighting at the level of the arena was 60 lux.

#### Cognitive Flexibility Test (Dynamic Strategy-Shifting Task)

##### Housing.

Prior to, and during operant training, mice were food deprived and weights were reduced and maintained at 90% of their free-feeding weights to enhance motivational salience of operant rewards. Training and testing were conducted from Monday-Saturday 1-2 times daily for the duration of experiments.

##### Apparatus.

Behavioural testing was conducted in modular operant chambers (Med Associates Inc., St. Albans, VT, USA). The Dynamic Strategy-Shifting Task (DSST) was conducted within six chambers (24.1cm x 20.3 x 18.4cm). Strawberry milk (Breaka, Queensland, Australia) was delivered as positive reinforcement via a motorised dipper mechanism by raising the dipper cup (∼0.06 cc) into the reward magazine. Reward magazines, nose-poke apertures and levers were all fitted with infrared head-entry detectors to record chamber interactions. The chambers also featured white-noise generators which were fitted externally to the back wall for the audio aspect of the task. All chambers were enclosed within sound-attenuating boxes (55.9cm x 55.9cm x 35.6cm), each fitted with a fan to provide ventilation and buffering from external noise, and an infrared camera (D-link Australia, North Ryde, NSW, Australia) for recording and observation. Protocols were written on MED-PC (Med Associates Inc. VT; S. Alexander) with a data collection resolution of 0.01s.

##### Dynamic Strategy-Shifting Task (DSST) test.

Instrumental training began with a “reward training” stage where the mice learned to collect rewards by making head entries (HEs) into the reward magazine to trigger the dipper mechanism. Mice were progressed to “self-initiation training” after completing the reward training stage. Here, they would learn how to ‘self-initiate a trial by making a HE into the centre nose-poke before making a HE into the magazine to collect a reward. The third and fourth stimulus-response training stages required the mice to first self-initiate a trial for the levers to be extended. Mice were then required to press either lever beside the magazine before making a HE into the magazine to collect a reward. The fourth stage differed only from the third stage by the introduction of further restrictions on the time to press a lever (limited hold) and to collect a reward. After completing these four stages, the mice would progress to the testing phase of the Dynamic strategy-shifting task.

The test phase of the DSST was delivered in two phases referred to as the primary and secondary exposures consisting of 5 and 7 stages, respectively. Details of stages are provided in **Error! Reference source not found.**A. Completion of the primary exposure was a requirement to progress to the secondary exposure. During primary exposure, mice were subjected to all spatial and non-spatial discrimination stages but not to non-spatial reversal stages until the secondary exposure. Criterion to pass each stage was 6 consecutive correct responses. If a mouse passed a stage within 30 minutes, it would progress to the next stage within that session. A session end was signalled if the mouse either failed to pass a stage within 30 minutes, made over 120 trials within one stage, had been inactive for over 5 minutes or the exceeded the maximum session time (90 minutes). Prior to commencement of test, mice were split into two groups with different starting target locations to control for any potential side bias. Number of trials to criteria (TTC) for each stage was the key behavioural outcome extracted from both training and testing phases.

##### Statistical Analyses.

Statistical analyses was conducted using the SPSS software package (ver.28, SPSS Inc. IL,USA) and GraphPad Prism (ver.9.3.1) with significant set to *p*<0.05. Only mice that had completed at least half of the stages in each exposure were included in TTC comparisons. Mice were given a maximum TTC of 350 if they never completed the stage or if they exceeded that amount for the purpose of conducting comparisons and reducing the effects of extreme outliers. Comparisons of TTC were conducted using Friedman tests and Wilcoxon signed rank tests with Dunn’s multiple comparisons test where appropriate, as windsorization resulted in non-gaussian distributions. Number of sessions was compared between genotypes using a Brown-Forsythe ANOVA.

## Acknowledgements

The work was funded by grants from the Australian Research Council (DP220100985 and DP230101750) to MP and ST. We thank Mikael Boden, Brad Balderson, and Rebecca Johnston for advice on the snRNA-seq analysis. We would also like to acknowledge Erica Mu and the histology team at the School of Biomedical Sciences (SBMS) for aid in Golgi-Cox staining and HREM processing, as well as Shaun Walters and the SBMS microscopy team. We appreciate the help provided by Virginia Nink for FACs sorting and Angelika Christ for processing nuclei for 10x Genomics sequencing. We thank Suzy Alexander for coding of the dynamic strategy shifting test and the Queensland Brain Institute (QBI) behaviour facility for training and providing the equipment for the behaviour tests. We acknowledge the supports from the Queensland NMR Network and the National Imaging Facility (a National Collaborative Research Infrastructure Strategy capability) for the operation of 16.4T MRI at the Centre for Advanced Imaging. Finally, we thank the QBI animal team for their exceptional care and housing of the animals. Arena schematics in Figure 7A, Supplementary 7A,G were created with BioRender.com

## Author contributions

Conceptualization, L.C., S.T., and M.P.; Methodology, L.C., S.T., B.M. M.A.-K., S.-J.M.; Investigation, L.C., B.M., M.A.-K., S-J.M., D.H., and A.P.; Formal analysis, L.C., B.M., M.A.-K., and S-J.M.; Writing – Original Draft, L.C.; Writing – Review & Editing, S.T., and M.P.; Supervision, N.K., T.B., L.H., S.T., and M.P.; Funding Acquisition, S.T., and M.P.; Project Administration, M.P.

---

**Supplementary 1.**
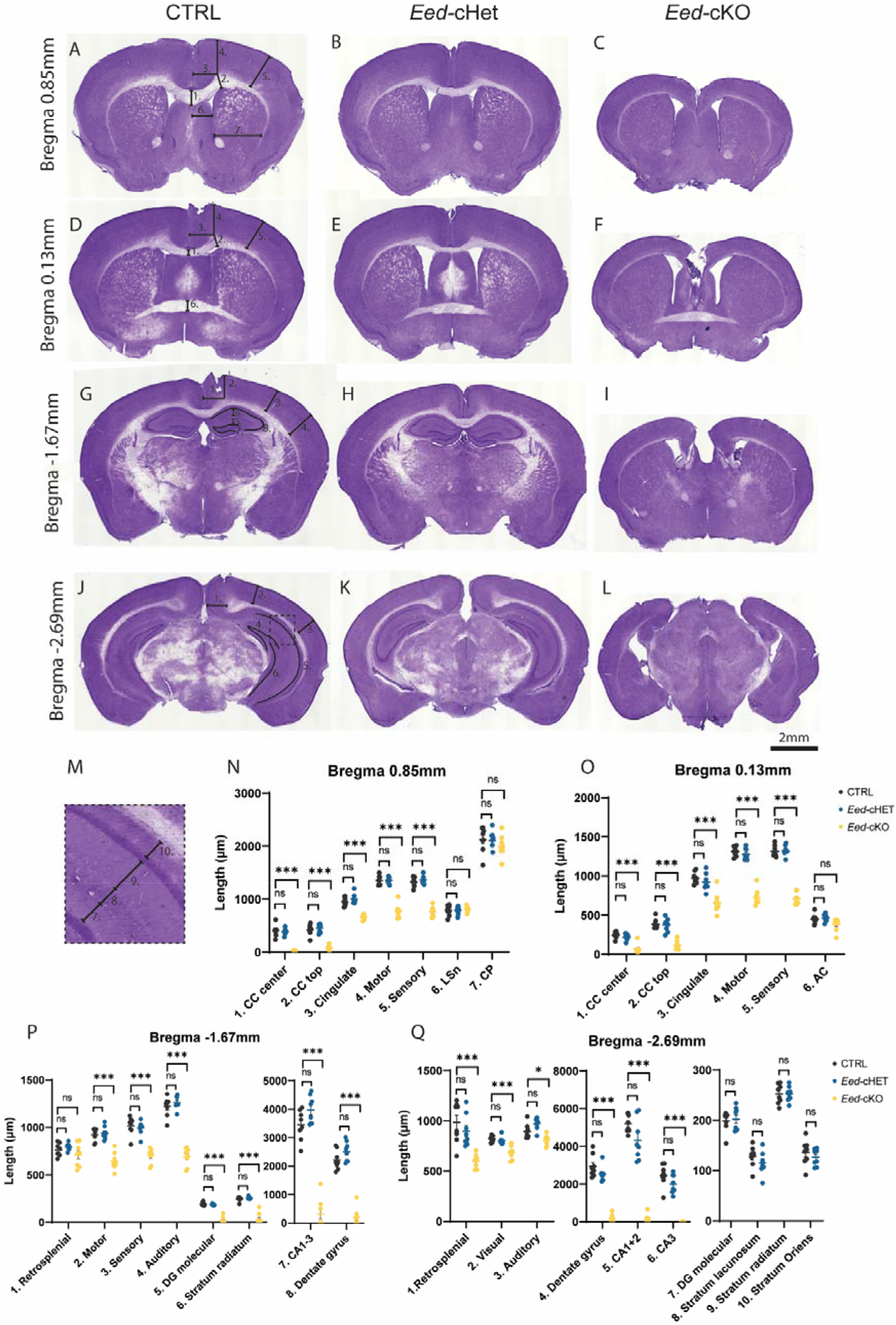
Morphometric analysis of *Eed-cHet* and *Eed-cKO* mice at Bregma 0.85mm, 0.13mm, −1.67mm, and −2.69mm. Hematoxylin-stained coronal sections of CTRL, *Eed-cHet*, and *Eed-cKO* brains at Bregma 0.85mm (A-C, N), Bregma 0.13mm (D-F, O), Bregma −1.67 (G-I, P), and Bregma −2.69 (J-L, Q). Annotations on (A, D, G, J, M) indicate how regions of interest were measured. CC = corpus callosum, LSn = lateral septal nucleus, AC = anterior commissure, HC = hippocampal commissure. *** = p<0.001, ns = not significant, one-way ANOVA. n = 8 CTRL, 8 *Eed-cHet*, 8 *Eed-cKO*.

**Supplementary 2.**
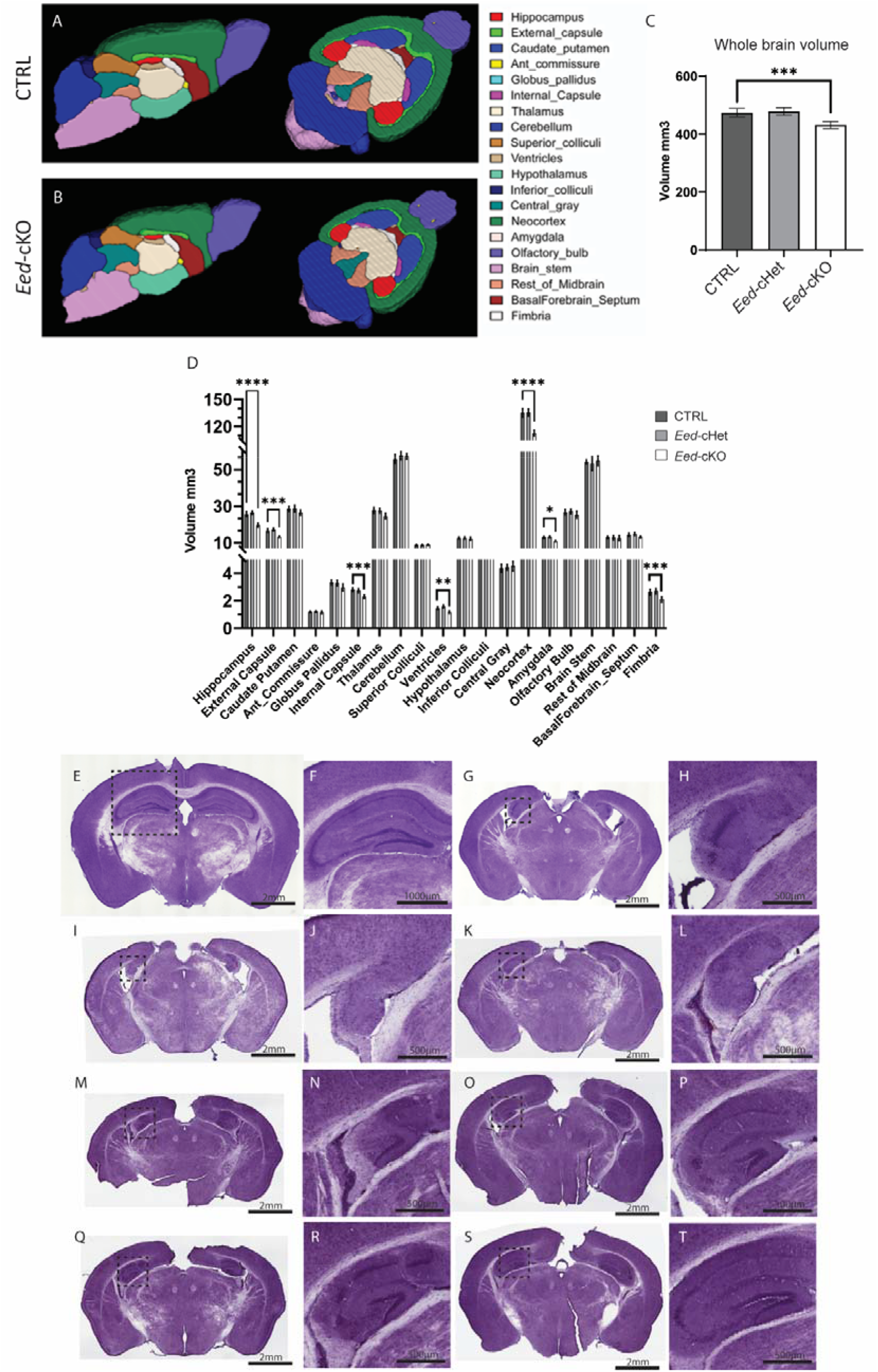
Structural MRI analysis and variation of Eed-cKO phenotype. (A-D) Structural MRI analysis of CTRL, *Eed-cHet*, and *Eed-cKO* mice. (A, B) Representative images of 3D reconstruction from MRI scans of control (A) and *Eed-cKO* (B) brains. (C) Measurement of whole brain volume. (D) Volume measurements of major brain regions. * = p<0.005, ** = p<0.001, ns = not significant. n = 7 CTRL, 8 *Eed-cHet*, 5 *Eed-cKO*. (E-T) Hematoxylin sections of one control (E, F) and 7 *Eed-cKO* biological replicates (G-T) demonstrates the variation of hippocampal phenotype in *Eed-cKO* brains. Panels (F, H, J, L, N, P, R, T) show zoomed-in images of the hippocampus of (E, G, I, K, M, O, Q, S), respectively. Panel H shows a *Eed-cKO* hippocampus with a structure resembling a CA region (yellow arrow), but no apparent dentate gyrus. Panel J and L show *Eed-cKO* brains which do not have any clear hippocampal structures. Panel N shows a *Eed-cKO* hippocampus with a structure resembling a dentate gyrus (orange arrow), but no apparent CA1-3 region. Panels P, R, and T show a *Eed-cKO* hippocampus which has both a CA region and dentate gyrus (yellow and orange arrow respectively), although they were clearly diminished compared to the control (B). Scale bar in (E, G, I, K, M, O, Q, S) = 2 mm. Scale bar in (B) = 100 µm. Scale bar in (F, H, J, L, N, P, R, T) = 50 µm.

**Supplementary 3.**
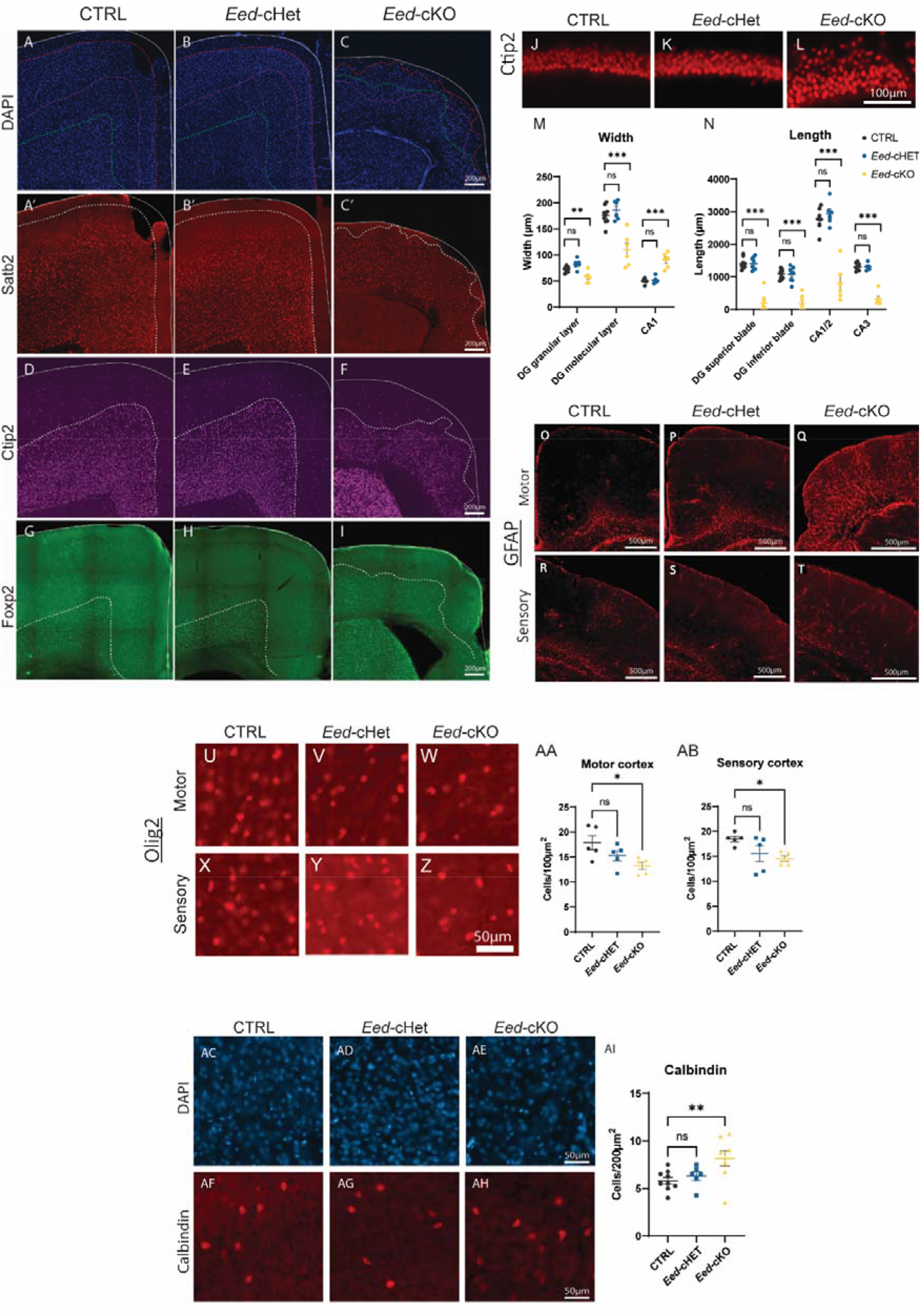
Additional immunofluorescent analysis of *Eed-cHet* and *Eed-cKO* brains. (A-I) Immunofluorescence staining of DAPI (A-C), Satb2 (A’-C’), Ctip2 (D-F), and Foxp2 (G-I) of the CTRL, *Eed-cHet*, and *Eed-cKO* motor/cingulate cortex. DAPI and Satb2 were co-stained, whereas Ctip2 and Foxp2 were stained from separate sections. Dashed lines represent the border between positive and negative layers for each marker. In CTRL and *Eed-cHet* samples, the border was an organised, straight line, whereas in *Eed-cKO* samples, the border was irregular and sometime ambiguous, indicating that cortical lamination was disorganised in *Eed-cKO* mice. Scale bar = 200 µm. (J-N) Supplementary data of immunofluorescence stains of Prox1, Calbindin, and Ctip2 in the hippocampus of CTRL, *Eed-cHet*, and *Eed-cKO* mice (presented in Figure 2). (J-K) Zoomed in image of the Ctip2+ CA1 region in CTRL, *Eed-cHet*, and *Eed-cKO* samples shows this region in greater detail. (M) The width of the dentate gyrus (DG) granular and molecular layers was measured, as well as the width of the CA1 region. (N) The length of the dentate gyrus superior and inferior blades, CA1/2, and CA3 regions were measured. (O-T) Immunofluorescence stains of GFAP in the motor (O-Q) and sensory (R-T) cortices of CTRL, *Eed-cHet*, and *Eed-cKO* mice. Compared to controls, *Eed-cKO* mice had increased expression of GFAP in the motor/cingulate cortex (Q), but not in the sensory cortex (T). (U-AB) Immunofluorescence staining of Olig2 in the motor (U-W) and sensory (X-Z) cortex of CTRL, *Eed-cHet*, and *Eed-cKO* mice. (AA-AB) Olig2+ cells were counted in the motor cortex (AA) and sensory cortex (AB). * = p<0.05, one-way ANOVA. n = 5 CTRL, 5 *Eed-cHet*, 5 *Eed-cKO*. (AC-AI) Immunofluorescence stain of DAPI and calbindin in the sensory cortex of CTRL, *Eed-cHet*, and *Eed-cKO* mice. Calbindin+ cells were counted in the middle layers of the sensory cortex (AI respectively). *** = p<0.001, ** = p<0.01, * = p<0.05, ns = not significant, one-way ANOVA. n = 10 CTRL, 8 *Eed-cHet*, 8 *Eed-cKO* (M-N); n = 5 CTRL, 5 *Eed-cHet*, 5 *Eed-cKO* (AA-AB); n = 7 CTRL, 6 *Eed-cHet*, 6 *Eed-cKO* (AI).

**Supplementary 4.**
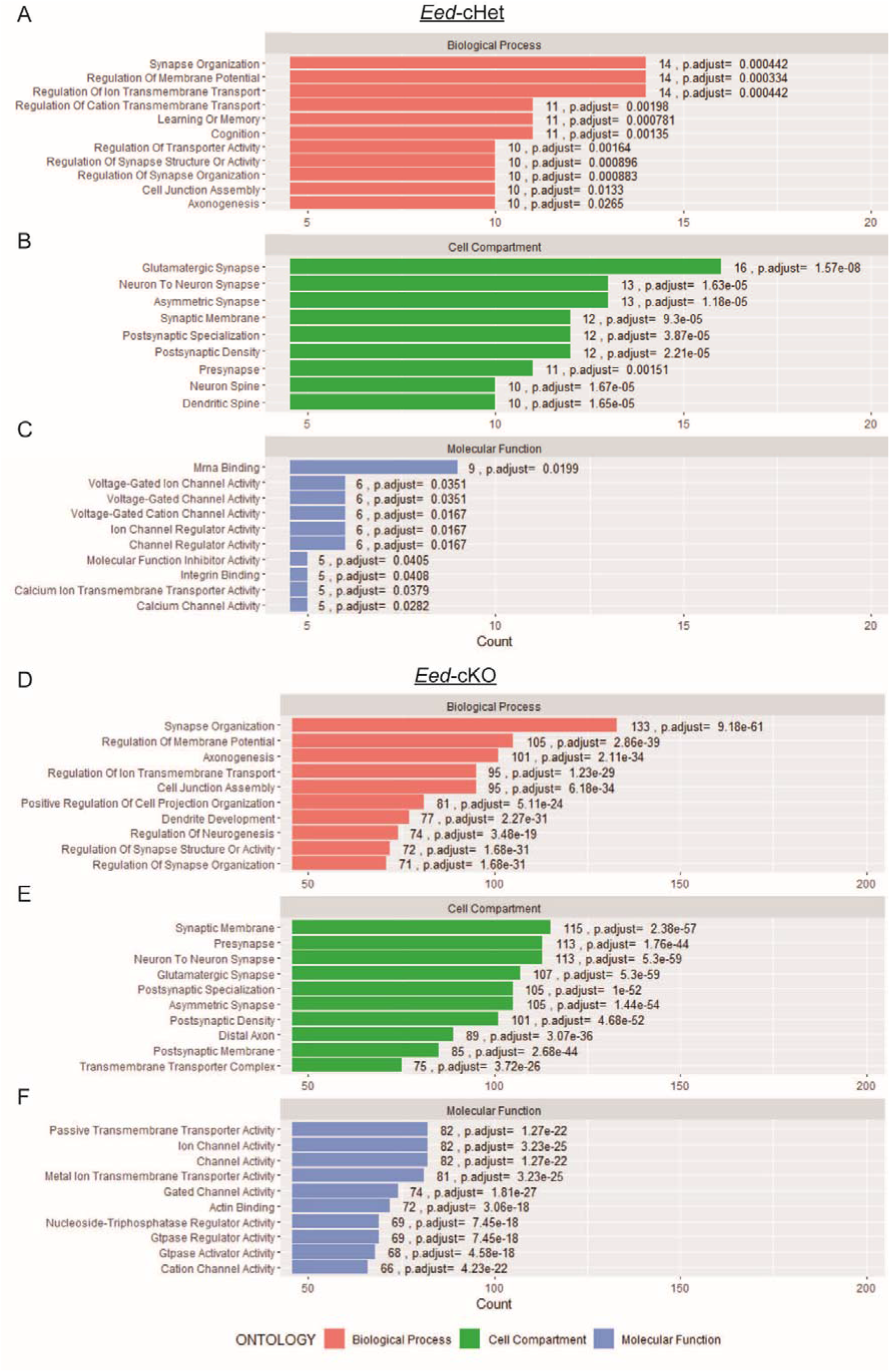
Gene ontology terms enriched in *Eed-cKO* mice. Top 10 enriched GO terms for *Eed-cHet* (A-C) and *Eed-cKO* (D-F) glutamatergic neurons. GO terms are categorized into biological processes (A, D), cell compartment (B, E), and molecular function (C, F). ‘Count’ refers to number of dysregulated genes that are associated with the GO term. Adjusted p-value is displayed next to each bar.

**Supplementary 5.**
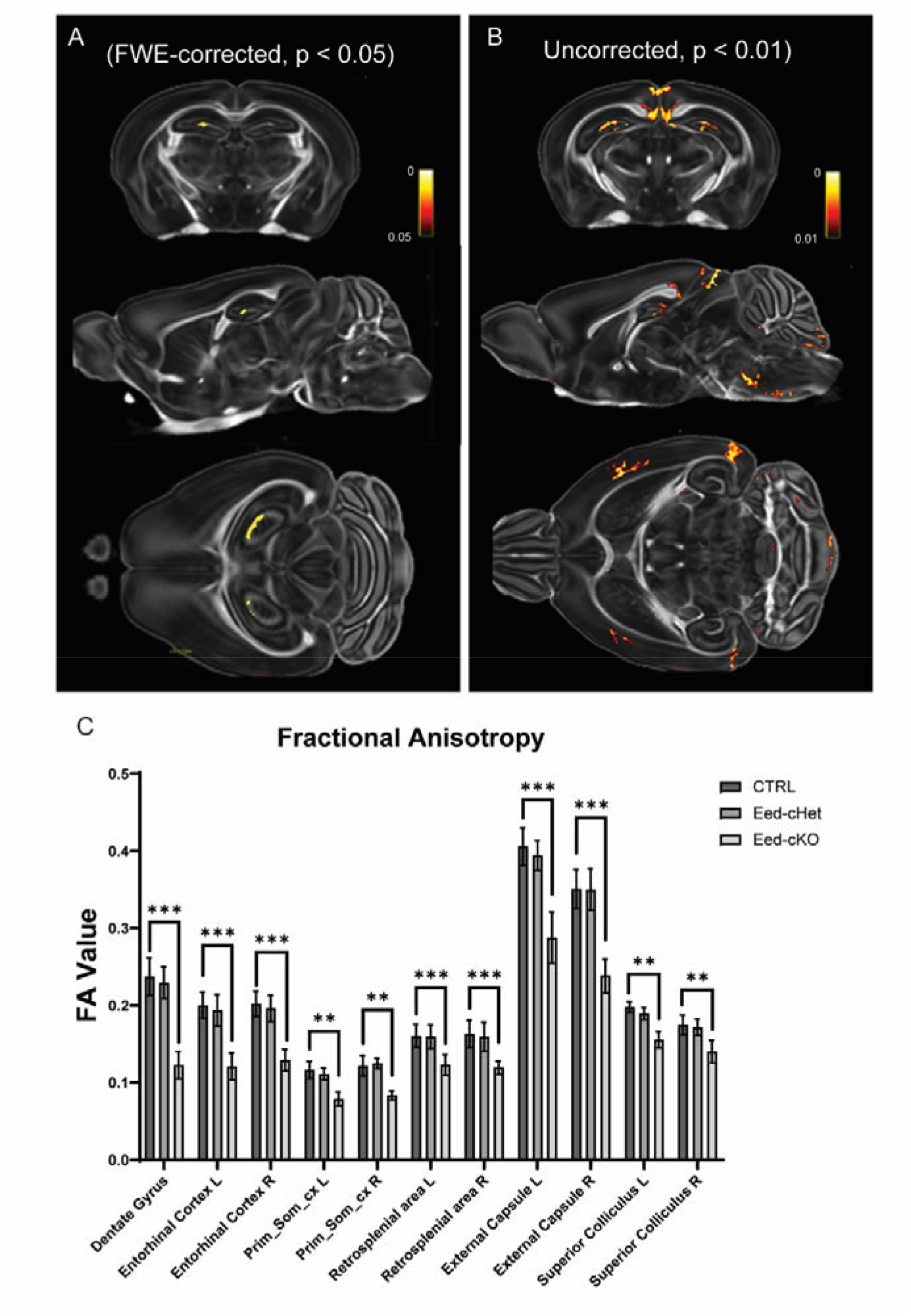
Voxel-based morphometry analysis of fractional anisotropy. (A) Map of significant changes in FA of *Eed-cKO* brains compared to control using a statistically stringent test corrected for family wise error (FWE) and p-value < 0.05. *Eed-cKO* mice had statistically reduced FA in the dentate gyrus. (B) Map of significant changes in FA of *Eed-cKO* brains compared to control using a less statistically stringent test, not corrected for FWE but with p-value < 0.01. (C) Graph showing each region where FA was statistically difference in *Eed-cKO* mice using the uncorrected test in (B). “L” and “R” refer to “left” and “right” respectively. “Prim_Som_cx” refers to the primary somatosensory cortex. Asterisks (*) indicate that p < 0.005. n = 7 CTRL, 8 *Eed-cHet*, 5 *Eed-cKO*.

**Supplementary 6.**
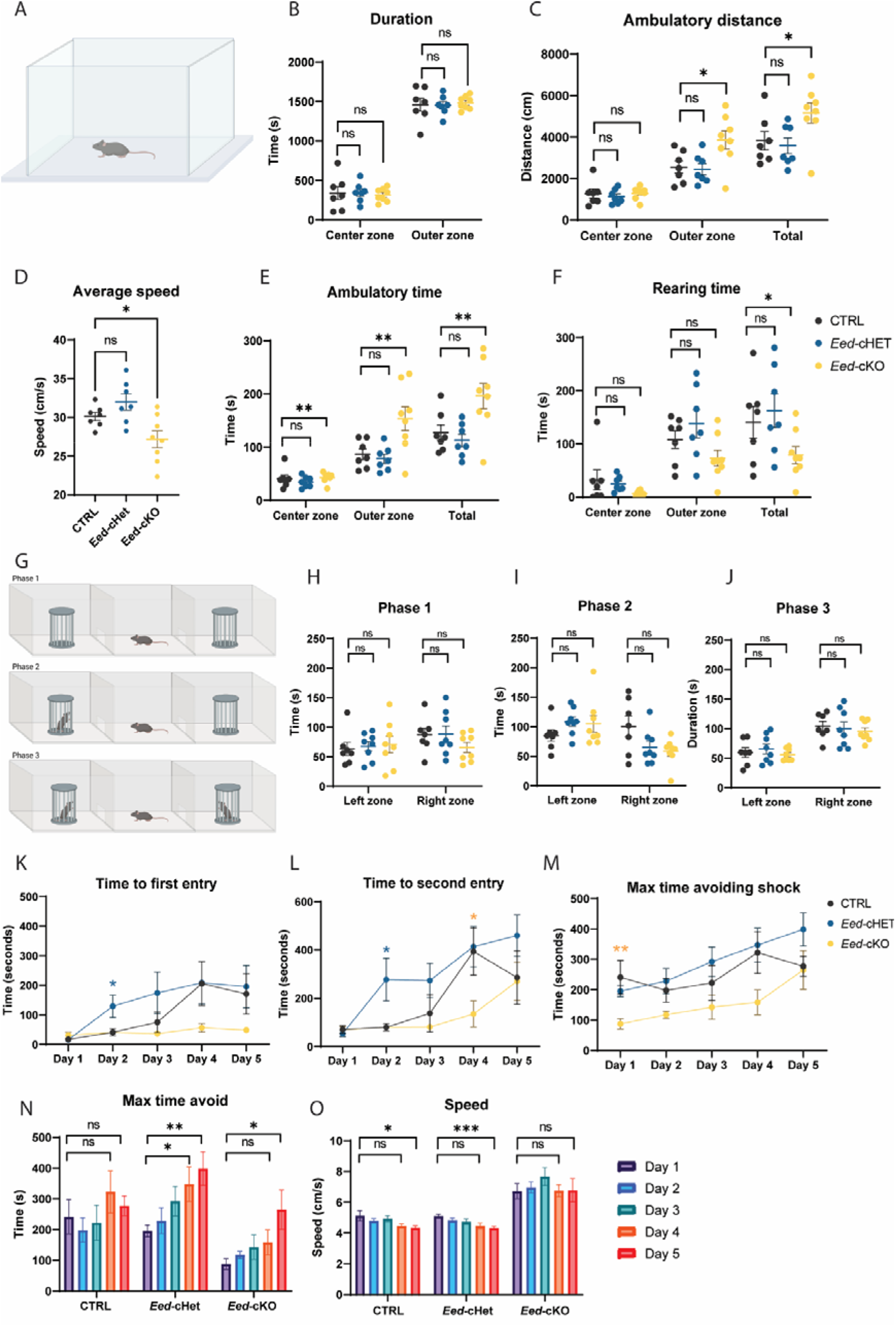
Open field maze, 3-chamber social interaction test, and additional data for the active place avoidance test. (A-F) Open field maze test. (A) Schematic of arena set up, which consists of an enclosed square arena. Duration in outer and inner zone (B), distance moved (C), and average speed (D) was measured. (E-F) Time spent performing activities including ambulatory time (E) and rearing time (F) was also measured. There was no significant difference in time spent on stereotypic movements, jumping or resting between genotypes (data not shown). (G-J) 3-chamber social interaction test. (G) Schematic of arena layout, which consists of 3 compartments, with cages in the outer compartments. The test consists of three phases in which the test mouse was acclimatised to the arena (phase 1), before being exposed to a novel mouse in the left cage (phase 2), and then a second novel mouse in the right cage (phase 3). Time spent next to the left and right cages (“left zone” and “right zone”, respectively) was measured. (K-O) Additional data for the active place avoidance test, as presented in Figure 7G-L. Intergenotype comparisons for time to first entry (K), time to second entry (L), and max time avoiding shock (M) are shown. Intragenotype comparisons for max time avoiding shock (N), and average speed (O) are shown. *** = p<0.001, * = p<0.05, ns = not significant, one-way ANOVA. n = 8 CTRL, 8 *Eed-cHet*, 8 *Eed-cKO*.

**Supplementary 7.**
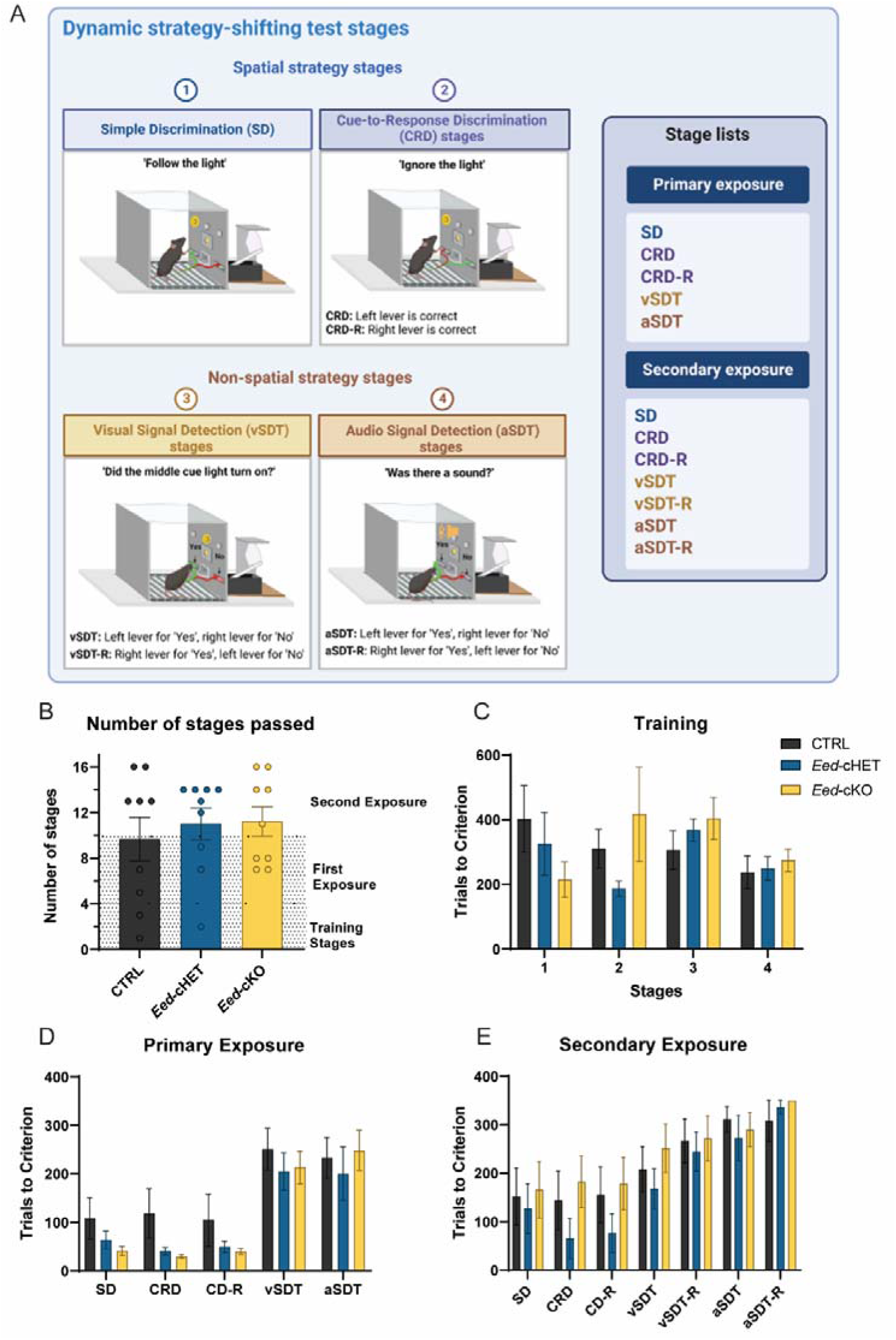
Dynamic strategy shifting test. (A) Schematic of arena and criteria for the primary and secondary exposure phases. (B) Number of stages passed. All three genotypes were able to complete all phases (training stage, first exposure, and second exposure). (C-E) Number of trials to criterion was measured for the training (C), primary exposure (D), or secondary exposure (E) phases. Statistical analysis was conducted with Brown-Forsythe ANOVA (B) or Friedman tests and Wilcoxon signed rank tests with Dunn’s multiple comparisons test where appropriate, as windsorization resulted in non-gaussian distributions (C-E). n= 9 CTRL, 9 *Eed-cHet*, 9 *Eed-cKO* (A); n= 7 CTRL, 8 *Eed-cHet*, 9 *Eed-cKO* (C-E).

**Figure 1.**
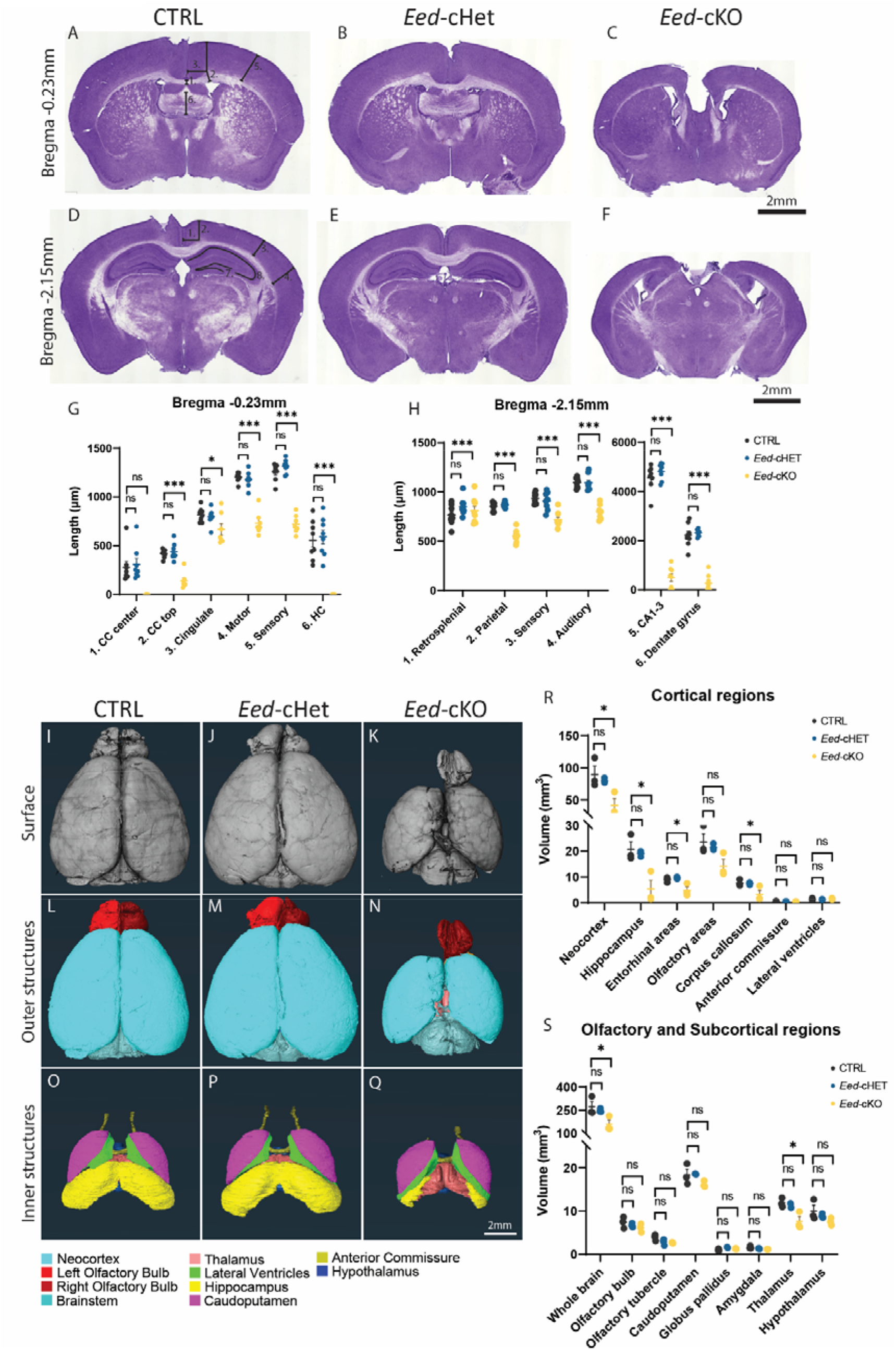
Loss of *Eed* results in a smaller cortex in the adult. (A-H) Hematoxylin-stained coronal sections at Bregma −0.23mm (A-C) and −2.15mm (D-F) of control CTRL, *Eed-cHet*, and *Eed-cKO* brains. (G, H) Regions of interest were measured as indicated by annotations on (A, D). CC = corpus callosum, HC = hippocampal commissure. *** = p<0.001, ns = not significant, one-way ANOVA. n = 8 CTRL, 8 *Eed-cHet*, 8 *Eed-cKO* (G-H). See also Supplementary 1 for analysis of additional Bregma sections. (I-S) HREM analysis of CTRL, *Eed-cHet*, and *Eed-cKO* brains. (I-Q) Representative images of HREM 3D reconstruction of the surface (A-C), outer structures (D-F), and inner structures (G-I) of CTRL (A, D, G), *Eed-cHet* (D, E, F), and *Eed-cKO* (G, H, I) brains. (J-K) The volume of major brain regions was measured. * = p<0.05, ns = not significant, one-way ANOVA. n = 3 CTRL, 3 *Eed-cHet*, 3 *Eed-cKO* (R-S).

**Table 1:** Expression of Prox1, Calbindin, and Ctip2 in the hippocampal regions of *Eed-cKO* mice.

|  | Prox1 | Calbindin | Ctip2 (CA1/2 region) |
| --- | --- | --- | --- |
| **Total examined** | 10 | 8 | 6 |
| **Expression in both hemispheres** | 3 | 3 | 4 |
| **Expression in one hemisphere** | 4 | 4 | 2 |
| **No expression** | 3 | 1 | 0 |

